# Cardiomyopathy Mutations Impact the Power Stroke of Human Cardiac Myosin

**DOI:** 10.1101/2020.12.08.416511

**Authors:** W. Tang, J. Ge, W.C. Unrath, R. Desetty, C. M. Yengo

## Abstract

Cardiac muscle contraction is driven by the molecular motor myosin that uses the energy from ATP hydrolysis to generate a power stroke when interacting with actin filaments, while it is unclear how this mechanism is impaired by mutations in myosin that can lead to heart failure. We have applied a Förster resonance energy transfer (FRET) strategy to investigate structural changes in the lever arm domain of human β-cardiac myosin subfragment 1 (M2β-S1). We exchanged the human ventricular regulatory light chain labeled at a single cysteine (V105C) with Alexa 488 onto M2β-S1, which served as a donor for Cy3ATP bound to the active site. We monitored the FRET signal during the actin-activated product release steps using transient kinetic stopped-flow measurements. We proposed that the fast phase measured with our FRET probes represents the structural change associated with rotation of the lever arm during the power stroke in M2β-S1. Our results demonstrated human cardiac muscle myosin has a slower power stroke compared with fast skeletal muscle myosin and myosin V. Measurements at different temperatures comparing the rate constants of the power stroke and phosphate release revealed that the power stroke occurs before phosphate release, and the two steps are tightly coupled. We speculate that the slower power stroke rate constant in cardiac myosin may correlate with the slower force development and/or unique thin filament activation properties in cardiac muscle. Additionally, we demonstrated that HCM (R723G) and DCM (F764L) associated mutations both reduced actin-activation of the power stroke in M2β-S1. We also demonstrate that both mutations decrease ensemble force development in the loaded in vitro motility assay. Thus, examining the structural kinetics of the power stroke in M2β-S1 has revealed critical mutation-associated defects in the myosin ATPase pathway, suggesting these measurements will be extremely important for establishing structure-based mechanisms of contractile dysfunction.

**Significance:** Mutations in human beta-cardiac myosin are known to cause various forms of heart disease, while it is unclear how the mutations lead to contractile dysfunction and pathogenic remodeling of the heart. In this study, we investigated two mutations with opposing phenotypes and examined their impact on ATPase cycle kinetics, structural changes associated with the myosin power stroke, and ability to slide actin filaments in the presence of load. We found that both mutations impair the myosin power stroke and other key kinetic steps as well as the ability to produce ensemble force. Thus, our results provide a structural basis for how mutations disrupt molecular level contractile dysfunction.

## Introduction

The myosin superfamily shares a highly conserved ATPase cycle that efficiently converts the energy from ATP hydrolysis into mechanical force (1, 2). One central question that continues to be debated in the field is how specific steps in the ATPase cycle are coupled to force generation. It is well understood that when myosin, containing the hydrolyzed products in its active site (ADP and Pi), binds to actin, it triggers structural rearrangements in the myosin motor, resulting in force generation (power stroke) and product release (3-5). However, the precise timing, as well as the sequence of events associated with the power stroke and product release, still remains elusive. Characterizing the structural and biochemical coupling between those critical steps is crucial to understand the myosin energy transduction mechanism.

The lever arm hypothesis suggests that force generation occurs when structural changes in the myosin active site are communicated to the lever arm domain, a long α-helix that extends from the motor domain that is stabilized by calmodulin-like light chains (3). The lever arm is known to undergo a ∼70° rotation that can produce force when myosin is tightly bound to actin. The lever arm rotation is thought to be coupled to actin-activated product release, especially phosphate release, which can be accelerated more than 1000-fold in the presence of actin (6). Different experimental techniques have been applied to study the structural and biochemical coupling in myosins, including single-molecule mechanics, X-ray crystallography, fluorescence spectroscopy, and cyro-electron microscopy (cryo-EM)(7-13). Additionally, FRET is a powerful approach to monitor the structural dynamics of lever arm rotation by engineering site-directed probes into the myosin molecule. FRET has been successfully utilized in several myosin isoforms including myosin V, skeletal myosin, and bovine cardiac myosin to investigate the structural kinetics of the lever arm rotation during the power stroke (14-19). The FRET results demonstrated that different myosin isoforms could display different biochemical properties of the power stroke, possibly tuning them for a specific physiological role. For example, the power stroke rate constants measured in myosin V and skeletal myosin are much faster than bovine cardiac myosin (14, 15, 17, 19). Therefore, it is important to perform FRET measurements of the power stroke in human β-cardiac myosin to better understand its force-generating properties, which will provide fundamental insights into understanding cardiac myosin motor performance in health and disease. Currently, more than 300 pathogenic single missense mutations have been reported clinically in the gene *MYH7*, encoding human β-cardiac myosin heavy chain (M2β)(20). Mutations in *MYH7* are commonly associated with hypertrophic cardiomyopathy (HCM) and dilated cardiomyopathy (DCM), while detailed molecular mechanisms of those mutations are still unclear. Understanding the molecular basis of disease-causing mutations will shed light on elucidating the structural and biochemical coupling between subdomains within the myosin molecule, as well as identifying potential therapeutic interventions. R723G and F764L are cardiomyopathy mutations both located in the converter domain, a region important for communication between the active site and lever arm, of M2β and clinically associated with HCM and DCM, respectively (21, 22). The converter domain is a hot spot for cardiomyopathy mutations in M2β and is also the load sensing region that undergoes most of the elastic distortion during the power stroke (23, 24). Therefore, mutations in the converter domain are hypothesized to be prone to alter the force-generating properties of M2β.

The biochemical impact of the F764L mutation in human M2β was well characterized together by previous work and our recent publication (25-29). Overall, F764L causes slightly reduced motor properties in M2β subfragment 1 (S1), containing the motor domain and entire light chain binding region, by slowing the actin-activated phosphate release and ADP release steps (25). Kinetic simulation results demonstrate that the duty ratio (fraction of ATPase cycle that myosin is in the force-generating states) is slightly decreased or not significantly altered by F764L in unloaded conditions (25, 26). The R723G mutation has been studied in human soleus muscle biopsy samples and isolated human cardiomyocytes (30-32). Interestingly, Kraft et al. (33) reported that the R723G mutation caused a reduced maximum force and similar Ca^2+^ sensitivity in isolated human cardiomyocytes, but higher maximum force and lower Ca^2+^ sensitivity in human soleus muscle fibers. Kawana et al. (34) characterized the R723G mutant using the M2β-shortS1 construct (containing the motor domain and essential light chain binding region). Interestingly, the HCM mutation R723G causes a decrease in maximum ATPase activity and intrinsic force but a 5-10% increase in sliding velocity. The impact of the R723G mutation on the detailed kinetics of key steps in the myosin ATPase was recently characterized by Vera et al. (35). They demonstrated that R723G increased the ADP release rate constant and weakened the ATP binding affinity, which resulted in a decrease in the duty ratio.

In current work, we successfully monitored the structural kinetics of the power stroke using site-specific FRET probes engineered into human M2β-S1. Our data suggest that human cardiac myosin has a relatively slow power stroke similar to that measured in bovine cardiac myosin, and the power stroke is tightly coupled to the phosphate release step. Our results clearly support a model in which the power stroke occurs before phosphate release. In addition, we examined the impact of the R723G mutation on the key steps of the myosin ATPase cycle with transient kinetic measurements and found that R723G causes an increase in the ADP release rate constant. We further examined the effect of R723G and F764L mutations on the power stroke. Despite the different clinical phenotypes observed in patients, both R723G and F764L cause a significant decrease in actin-activation of the power stroke. We also demonstrated a reduction in the ensemble force generated by R723G and F764L in the loaded in vitro motility assay. Our results provide important insights into the molecular mechanisms of cardiomyopathy mutations and the structural basis of the myosin power stroke.

## Materials and Methods

### Reagents

Cy3ATP and Cy3ADP were purchased from Jena Bioscience (1 mM stock). Alexa Fluor 488 C5 Maleimide powder was purchased from Invitrogen and dissolved in DMSO. ATP and ADP were prepared from powder (MilliporeSigma) and concentrations were determined by absorbance at 259 nm (ε_259_ = 15,400 M^-1^cm^-1^). The fluorescently labeled phosphate binding protein (MDCC-PDB) was prepared as described (36). MOPS 20 buffer (10 mM MOPS, 20 mM KCl, 1 mM EGTA, 1 mM MgCl2, 1 mM DTT, pH 7.0) was used for all solution experiments.

### Protein construction, expression, and purification

We cloned the cDNA of M2β-S1 (841 aa) into a pshuttle vector which contains a C-terminal Avi tag and N-terminal FLAG tag (DYKDDDDK). The R723G and F764L point mutations were introduced using Quikchange site-directed mutagenesis (Stratagene). Recombinant adenovirus was produced and used to infect C_2_C_12_ cells as previously described (25, 37). Cells were harvested 7-10 days after infection and purified with FLAG affinity chromatography. Endogenous mouse RLC was removed from M2β-S1 using an on-column stripping method described previously (38). Briefly, S1 was bound to the FLAG column, by first flowing through cell lysate and washing the resin with wash buffer (10 mM Tris, pH 7.5, 200 mM KCl, 1 mM EGTA, 1 mM EDTA, 2 mM MgCl_2_, 2 mM ATP, 1 mM DTT, 0.01 mg/ml aprotinin, 0.01 mg/ml leupeptin, and 1 mM PMSF), and then incubated with stripping buffer (20 mM Tris, pH 7.5, 0.5% Triton X-100, 200 mM KCl, 5 mM CDTA, 2 mM ATP, 1 mM DTT, 0.01 mg/ml aprotinin, 0.01 mg/ml leupeptin, and 1 mM PMSF) for 1h, followed by incubation with 20 µM Alexa 488-labeled human RLC (A488RLC) in wash buffer to allow the exchange to occur. The exchanged M2β-S1 was washed again to remove exogenous A488RLC and eluted with FLAG peptide. About 0.5-1 mg of protein was obtained per 20 plates of C_2_C_12_ cells. M2β-S1 used for the *in vitro* motility assays was biotinylated with BirA (10 µg/ml, Avidity) as described, followed by ammonium sulfate precipitation (25). An N-terminally His tagged version of the human ventricular cardiac regulatory light chain (hRCL) with a single reactive cysteine at position 105 was expressed using pET15b vector in *E. coli* cells. The cell lysate was collected and purified with Talon-Affinity chromatography (His tag). The purified hRLC was labeled with 5 molar excess Alexa Fluor 488 overnight at 4 °C, and dialyzed into wash buffer to remove excess dye. The labeled A488 RLC was snap frozen in liquid nitrogen and kept at -80 °C until use. Actin was purified using acetone powder from rabbit skeletal muscle (39), and labeled with pyrene iodoacetamide when needed (40). All proteins were dialyzed in MOPS 20 buffer overnight before experiments.

### Steady-state ATPase measurements

0.1 µM M2β-S1 in the presence of varying actin concentrations (0, 5, 10, 20, 40, 60 µM) was examined in actin-activated ATPase experiments using a NADH-coupled ATP regenerating system as previously described (41, 42). Data were acquired for 200 seconds (0.2 s intervals) at 25 °C using an Applied Photophysics (Surrey, UK) stopped-flow. The ATPase rate was plot as a function of actin concentration and fit to the Michaelis-Menten equation to determine the *k*_cat_ and *K*_ATPase_. Results of wild type (WT) and two mutants were reported as the average of 3-6 protein preparations. Statistical analysis was done with unpaired student’s *t*-tests to determine the impact of A488RLC exchange and compare each mutant with WT.

### *In vitro* motility

The actin sliding velocity was measured as previously described (25). 1% nitrocellulose coated coverslips were incubated with streptavidin (0.1 mg/ml) and blocked with BSA (2 mg/ml). Biotinylated M2β-S1 WT or R723G or F764L was attached to the surface at 0.24 µM loading concentration. Sheared unlabeled actin (2 µM) followed by 2 mM ATP were added to block the dead myosin heads on the surface. The activation buffer containing MOPS 20 buffer, 0.35% methylcellulose, 0.45 mM phosphoenolpyruvate, 45 units/ml pyruvate kinase, 0.1 mg/ml glucose oxidase, 5 mg/ml glucose, 0.018 mg/ml catalase, and 2 mM ATP was added right before video acquisition. Alexa 488 or Rhodamine 555 phalloidin-labeled actin was visualized with a Nikon TE2000 fluorescence microscope. Videos were collected for 2 min at a 1 s frame rate. The velocities were either manually analyzed by tracking actin filaments using MTrackJ in ImageJ (43) or processed by the program FAST (44).

In the loaded *in vitro* motility assay, the M2β-S1 motor-dead mutant (E466A) protein was mixed with WT, R723G or F764L before added to the coverslip, and the total amount of M2β-S1 loaded remained constant at 0.6 µM. Relative velocities were normalized to the velocity at 0% motor-dead. Unpaired student’s *t*-tests were performed to compare each mutant with WT and the impact of A488RLC exchange.

### Transient kinetic measurements

An applied Photophysics stopped-flow equipped with an excitation monochromator, 1.2 ms dead-time, and a 9.3 nm band pass was used for all experiments. Tryptophan fluorescence was excited at 290 nm and monitored with a 320 nm long pass emission filter. The *mant* fluorescence was examined with 290 nm excitation and a 395 nm long pass emission filter. Pyrene actin fluorescence was monitored with 365 nm excitation and a 395 nm long pass emission filter. MDCC-PBP was monitored with a 380 nm excitation and a 425 nm long pass emission filter. Fluorescence transients were fit with the stopped-flow program or GraphPad Prism.

### FRET measurements

The steady-state FRET was measured by exciting the sample at 460 nm and measuring the fluorescence emission as a function of wavelength (480-650 nm) using a PTI spectrofluorimeter equipped with excitation and emission monochromators and Peltier temperature control cuvette holder. Stopped-flow FRET was examined by monitoring the change in donor fluorescence (Alexa 488) using an excitation wavelength of 470 nm and measuring the emission with an interference filter (500-525nm), which eliminated background fluorescence from Cy3ATP or Cy3ADP (absorbance peak of 550 nm and emission peak of 570 nm). The steady-state FRET efficiency (*E*) was calculated by examining donor quenching using the following equation (45, 46),

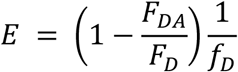

where *F*_DA_ is the donor fluorescence intensity in the presence of acceptor, *F*_D_ the donor fluorescence intensity in the absence of acceptor, and *f*_D_ is the fractional labeling with donor. The fraction of M2β-S1 with Cy3 nucleotide bound (*f*_D_) was determined from the *K*_D_, estimated from the stopped flow measurements (Figure 1). The distance (*r*) between the donor and acceptor was calculated based on the equation below (46),

**Figure 1.**
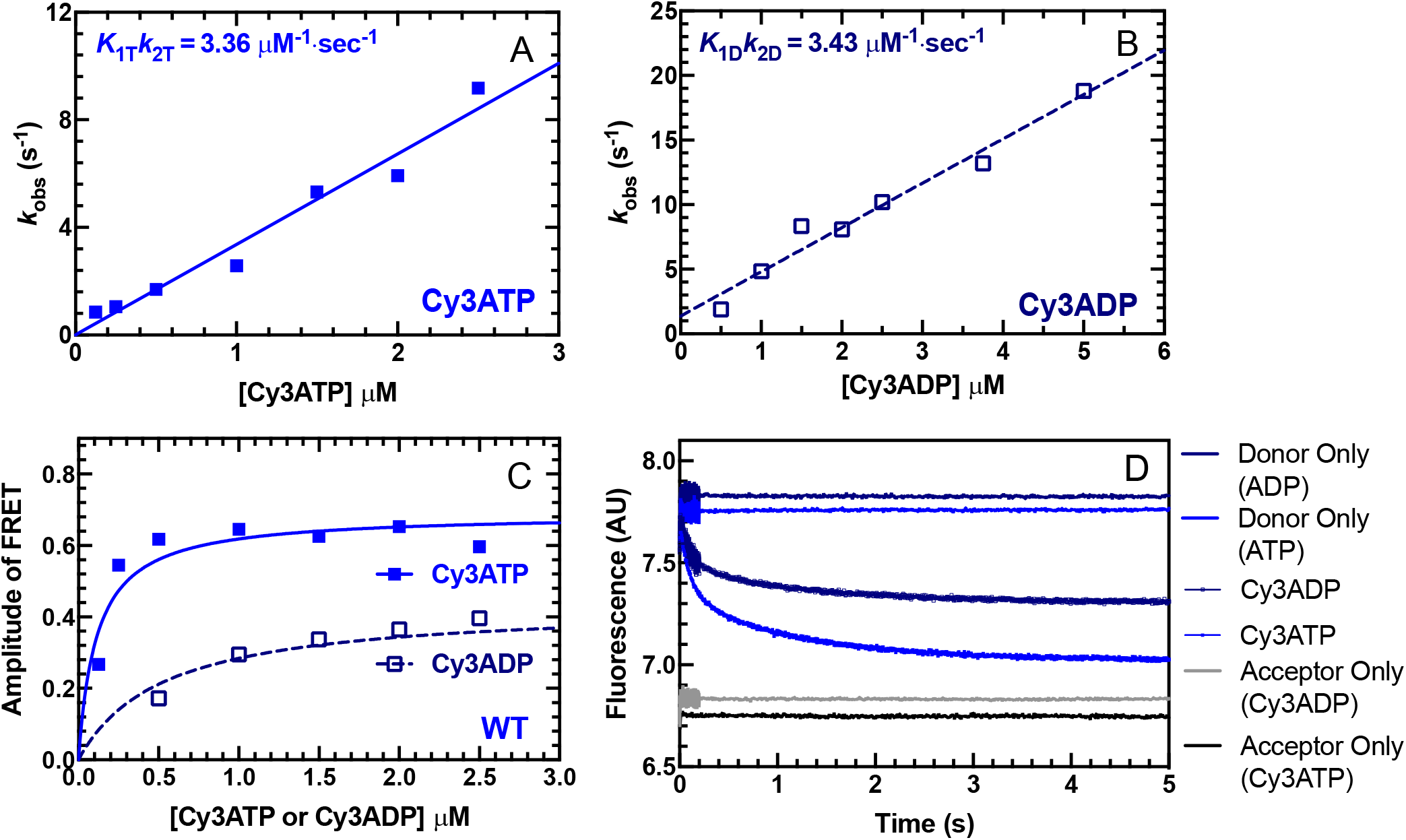
Cy3-labeled nucleotide binding to M2β-S1. The FRET change observed upon Cy3ATP or Cy3ADP binding to M2β-S1 was monitored by mixing 1 µM M2β-S1 A488RLC with varying concentrations of fluorescent nucleotide and monitoring the decrease in donor fluorescence. Fluorescence transients were best fit by a double exponential function. **(A)** The observed fast phase rate constant for Cy3ATP binding to M2β-S1 was linearly dependent on Cy3ATP concentration which allowed determination of the second-order binding constant. **(B)** Similarly, the observed fast phase rate constant for Cy3ADP binding to M2β-S1 was also linearly dependent on Cy3ADP concentration. **(C)** The amplitude of the FRET signal in the Cy3ATP and Cy3ADP experiments was plotted as a function of nucleotide concentration and was fit to a hyperbolic function to compare the relative amplitudes of the FRET signal (Cy3ATP, *A*_Max_ = 0.69 ± 0.04; Cy3ADP, *A*_Max_ = 0.43 ± 0.04). **(D)** Representative fluorescent transients from the Cy3ATP and Cy3ADP binding experiments fit to a double exponential function (Cy3ATP, *k*_Fast_ = 8.8 ± 0.1 s^-1^, *A*_Fast_ = 0.52; Cy3ADP, *k*_Fast_ = 8.4 ± 0.1 s^-1^, *A*_Fast_ = 0.54). No change in fluorescence was observed with donor only and acceptor only controls.

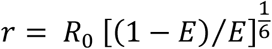

where the Förster distance (*R*_0_), the distance at which energy transfer is 50% efficient, was determined to be 67Å (17). Unpaired student’s *t*-tests were performed to compare differences, between the mutants and WT, in the FRET distance in each nucleotide state.

### Kinetic modeling and duty ratio

The fluorescence transients of the power stroke and phosphate release were fit to kinetic models as shown in Scheme 1 using Kintek Explorer (47, 48). The forward and reverse rates constants used for modeling the steady-state ATPase and specific transient kinetics steps are listed in Table S3.The duty ratio was estimated using the following equation (49):

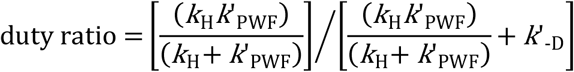

where *k*_H_ is the maximum rate of ATP hydrolysis measured by intrinsic tryptophan fluorescence, *k*’_PWF_ is the maximum rate of the fast power stroke rate constant, *k*’_-D_ is the rate of ADP release from actomyosin monitored with pyrene actin fluorescence.

## Results

### Purification and preparation of cardiac myosin constructs

M2β-S1 proteins were expressed and purified in C_2_C_12_ cells as previously described (25). Purified M2β-S1 contained the S1 fragment of human cardiac myosin heavy chain (amino acids 1-841) and associated mouse essential and regulatory light chains, which are endogenously expressed in C_2_C_12_ cells. Our previous work determined the sequences of the two mouse light chains using LC-MS/MS and demonstrated they are highly homologous to the human isoforms (41). To examine the structural kinetics of lever arm rotation by FRET, we exchanged the mouse RLC with Alexa 488-labeled human RLC (A488RLC, labeled at V105C) during the M2β-S1 purification process (see Methods for details). For all M2β-S1 WT, R723G, and F764L constructs, non-exchanged M2β-S1 protein was prepared in parallel with the exchanged protein and served as a control in all FRET experiments. Human RLC labeling and exchange efficiencies were determined by examining Alexa 488 concentration by absorbance and utilizing fluorescence gel quantitation methods (Figure S1). Briefly, M2β-S1 A488RLC complexed with actin was pelleted by ultracentrifugation and then released with ATP, the amount of A488RLC in the supernatant was quantified using an in-gel standard curve with an A488RLC standard. The total concentration of M2β-S1 A488RLC in the supernatant was determined by Bradford. We found that the stoichiometry of A488RLC to M2β-S1 was close to 1:1 (1:1 ± 0.2, for WT M2β-S1 A488RLC, *N* = 3).

### Steady-state motor properties of RLC-exchanged constructs

We first examined the impact of exchanging A488RLC onto M2β-S1 with steady-state ATPase and *in vitro* motility assays. The actin-activated ATPase activity was plotted as a function of actin concentration and fit to a Michaelis-Menten equation to determine the maximum rate (*k*_cat_) and actin concentration as which ATPase in one-half maximal (*K*_ATPase_) (Figure S2 and Table S1). The maximum rate was slightly lower in M2β-S1 A488RLC (4.4 ± 0.6 s^-1^) compared with non-exchanged controls (5.4 ± 1.3 s^-1^), but the difference was not significant (Figure S2A and Table S1). The average sliding velocity was examined in the *in vitro* motility assay, and velocity values were analyzed by the program FAST (44). The M2β-S1 A488RLC displays a slightly slower sliding velocity compared with the M2β-S1 (Figure S2B and Table S1). We hypothesized the difference could be due to the different RLC isoforms. We further examined the actin-activated ATPase and sliding velocity of M2β-S1 exchanged with unlabeled human RLC (V105C), and we found a slight decrease in actin sliding velocity and no change in maximum ATPase rate (5.3 ± 0.9 s^-1^)(Table S1), suggesting the presence of the human RLC slightly changes the mechanochemistry of M2β-S1. Overall, we conclude that the exchange of labeled human A488RLC onto M2β-S1 only causes minor changes to the steady-state motor properties of M2β-S1.

### Steady-state and transient FRET measurements

Previously, Rohde et al. (17) demonstrated a method of measuring lever arm rotation using Cy3-labeled nucleotides and A488RLC in bovine cardiac myosin. In the current work, we used a similar approach, and successfully monitored the FRET signal associated with lever arm rotation using human M2β-S1 A488RLC. Our transient kinetic results are interpreted in the context of Scheme 1, which we have utilized in previous studies (25). We observed different FRET efficiencies between Cy3ATP (0.48 ± 0.01) and Cy3ADP (0.40 ± 0.02) when examining 0.5 µM M2β-S1 A488RLC in the presence of 1 µM Cy3 labeled nucleotide under equilibrium conditions (corrected for fraction bound, see Methods) (Figure S3 and Table 1). We calculated the average distance change between donor and acceptor in the Cy3ATP and Cy3ADP conditions (ΔFRET = 3.8 ± 1.0 Å), which is similar to the distance change expected from structural modeling in the pre- and post-power stroke states (estimated distances of 84.3 and 86.9 Å, respectively)(Figure S4 and Table 1). Direct binding of Cy3ADP or Cy3ATP was monitored by mixing 1 µM M2β-S1 A488RLC with varying concentrations of Cy3ATP (0.1 to 2.5 µM) or Cy3ADP (0.5 to 5 µM) (Figure 1A&B) in the stopped-flow apparatus. The second-order binding constants were similar to published values for ATP and ADP (Cy3ATP, 3.36 ± 0.16 µM^-1^·s^-1^; Cy3ADP, 3.43 ± 0.28 µM^-1^·s^-1^), respectively (25, 50). Consistent with steady-state FRET, the maximum amplitude of the FRET change was larger with Cy3ATP (0.69 ± 0.04) than Cy3ADP (0.43 ± 0.04) (Figure 1C), indicating our FRET method monitors average lever arm position as a function of nucleotide-state.

**Table 1.** Steady-state FRET efficiency in different nucleotide states.

| FRET Parameter, $N = 6$, ±SEM | WT | R723G | F764L |
| --- | --- | --- | --- |
| Cy3ATP FRET Efficiency | $0.48 \pm 0.01$ | $0.45 \pm 0.01$ | $0.48 \pm 0.01$ |
| Cy3ADP FRET Efficiency | $0.40 \pm 0.02$ | $0.36 \pm 0.02$ | $0.35 \pm 0.02$ |
| Cy3ATP FRET Distance ($\text{\AA}$) | $67.9 \pm 0.5$ | $69.3 \pm 0.5$ | $67.9 \pm 0.5$ |
| Cy3ADP FRET Distance ($\text{\AA}$) | $71.7 \pm 1.0$ | $73.7 \pm 1.1$ | $74.5 \pm 1.1$ |
| ΔFRET ($\text{\AA}$)* | $3.8 \pm 1.0$ | $4.4 \pm 1.1$ | $6.6 \pm 1.1$ |

\*Difference between the distances determined in the presence Cy3ATP and Cy3ADP.

### Structural kinetics of lever arm rotation during the power stroke

A sequential mix stopped-flow apparatus was used to measure the kinetics of lever arm rotation during the power stroke. 0.25 µM M2β-S1 A488RLC was first mixed with 0.2 µM Cy3ATP, aged for 10 s to allow ATP hydrolysis to occur, and then mixed with varying concentrations of actin (5-30 µM) (Figure 2A). The traces (Figure 2B) were best fit to a double exponential function. The rate constants of the two phases (fast and slow) at different actin concentrations were fit to a hyperbolic function. We determined the fast phase of the lever arm rotation has a maximum rate of 17.2 ± 4.8 s^-1^ (Table 2). The slow phase was also actin-dependent but did not fit well to a hyperbolic function (rate constant of 1.5 s^-1^ at 30 µM actin). To further characterize the fast and slow phases of the power stroke, we repeated the power stroke measurements with 30 µM actin at different temperatures and plotted the rate constants together with temperature-dependent ATPase measurements (Figure 2C). We found the fast phase of the power stroke was faster than the ATPase values while the slow phase was slower than ATPase at all temperatures examined. The Eyring plot comparing the fast and slow phases with ATPase values (Figure 2D) showed that the three rate constants share a similar temperature dependence (slope, *k*_Fast_, -17.03 ± 0.76; *k*_Slow_, -18.25 ± 1.85; ATPase, -15.53 ± 0.51).

**Table 2.** Summary of steady-state and transient kinetic results comparing WT and mutants.

|  | WT | R723G | F764L |
| --- | --- | --- | --- |
| **Steady-State ATPase Values, (± SEM), $N = 3-6$** |  |  |  |
| $v_0$ ($s^{-1}$) | $0.02 \pm 0.01$ | $0.03 \pm 0.01$ | $0.02 \pm 0.01$ |
| $k_{cat}$ ($s^{-1}$) | $8.3 \pm 1.1$ | $6.1 \pm 1.5$ | $6.4 \pm 1.4$ |
| $K_{ATPase}$ ($\mu M$) | $78 \pm 16$ | $60 \pm 26$ | $70 \pm 25$ |
| ***In vitro* motility value, (± SEM), $N = 90$** |  |  |  |
| Mean Velocity (nm/s) | $1591 \pm 16$ | $1699 \pm 17^*$ | $1425 \pm 16^*$ |
| **Rate/Equilibrium Constants (± SEM)** | **$f_{WT}$** | **R723G** | **$f_{F764L}$** |
| **<sup>a</sup>ATP Binding/Hydrolysis (Myosin), $N = 2$** |  |  |  |
| $K_{0.5}$ ($\mu M$) | $48 \pm 6$ | $48 \pm 9$ | $79 \pm 6$ |
| $K_{1T}k_{+2T}$ ($\mu M^{-1} \cdot s^{-1}$) | $2.8 \pm 0.1$ | $3.9 \pm 0.1$ | $1.9 \pm 0.1$ |
| $k_{+H} + k_{-H}$ ($s^{-1}$) | $182 \pm 6$ | $231 \pm 12$ | $166 \pm 6$ |
| **<sup>b</sup>ATP Binding (Actomyosin), $N = 3$** |  |  |  |
| $K'_{1T}k'_{+2T}$ ($\mu M^{-1} \cdot s^{-1}$) | $9.6 \pm 0.2$ | $8.2 \pm 0.1$ | $9.4 \pm 0.2$ |
| $K_{0.5}$ ($\mu M$) | $31 \pm 9$ | $42 \pm 22$ | $24 \pm 8$ |
| $k'_{+2T}$ ($s^{-1}$) | $512 \pm 47$ | $668 \pm 121$ | $409 \pm 42$ |
| **<sup>c</sup>Actin-activated Pi Release at 30 $\mu M$** |  |  |  |
| $k_{+Pi}$ (fast, $s^{-1}$) | $20.7 \pm 0.9$ | ND | $16.4 \pm 1.7$ |
| $k_{+Pi}$ (slow, $s^{-1}$) | $1.86 \pm 0.02$ | ND | $1.68 \pm 0.01$ |
| **<sup>d</sup>Power Stroke, $N = 3-4$** |  |  |  |
| $k_{+PWF}$ (fast phase, maximum rate, $s^{-1}$) | $17.2 \pm 4.8$ | $10.8 \pm 4.1$ | $8.7 \pm 4.5$ |
| $K_{0.5, fast}$ ($\mu M$) | $48 \pm 33$ | $29 \pm 19$ | $29 \pm 26$ |
| $k_{+PWS}$ (slow phase, at 30 $\mu M$ Actin, $s^{-1}$) | $1.5 \pm 0.3$ | $0.9 \pm 0.2$ | $0.8 \pm 0.3$ |
| **<sup>e</sup>Actomyosin ADP Release,** |  |  |  |
| $^e k'_{+D}$ ($s^{-1}$), $N = 3-5$ | $332 \pm 57$ | $380 \pm 55$ | $290 \pm 46$ |
| $^b k'_{+D}$ ($s^{-1}$), $N = 3$ | $174 \pm 12$ | $218 \pm 15^*$ | $132 \pm 16^*$ |

<sup>a</sup> Intrinsic tryptophan fluorescence
<sup>b</sup> Pyrene actin
<sup>c</sup> MDCC-PBP fluorescence
<sup>d</sup> FRET
<sup>e</sup> *mant*-ADP fluorescence
<sup>f</sup> Values acquired from Tang et al.(25) except for the power stroke parameters
<sup>\*</sup>, $p < 0.05$

**Figure 2.**
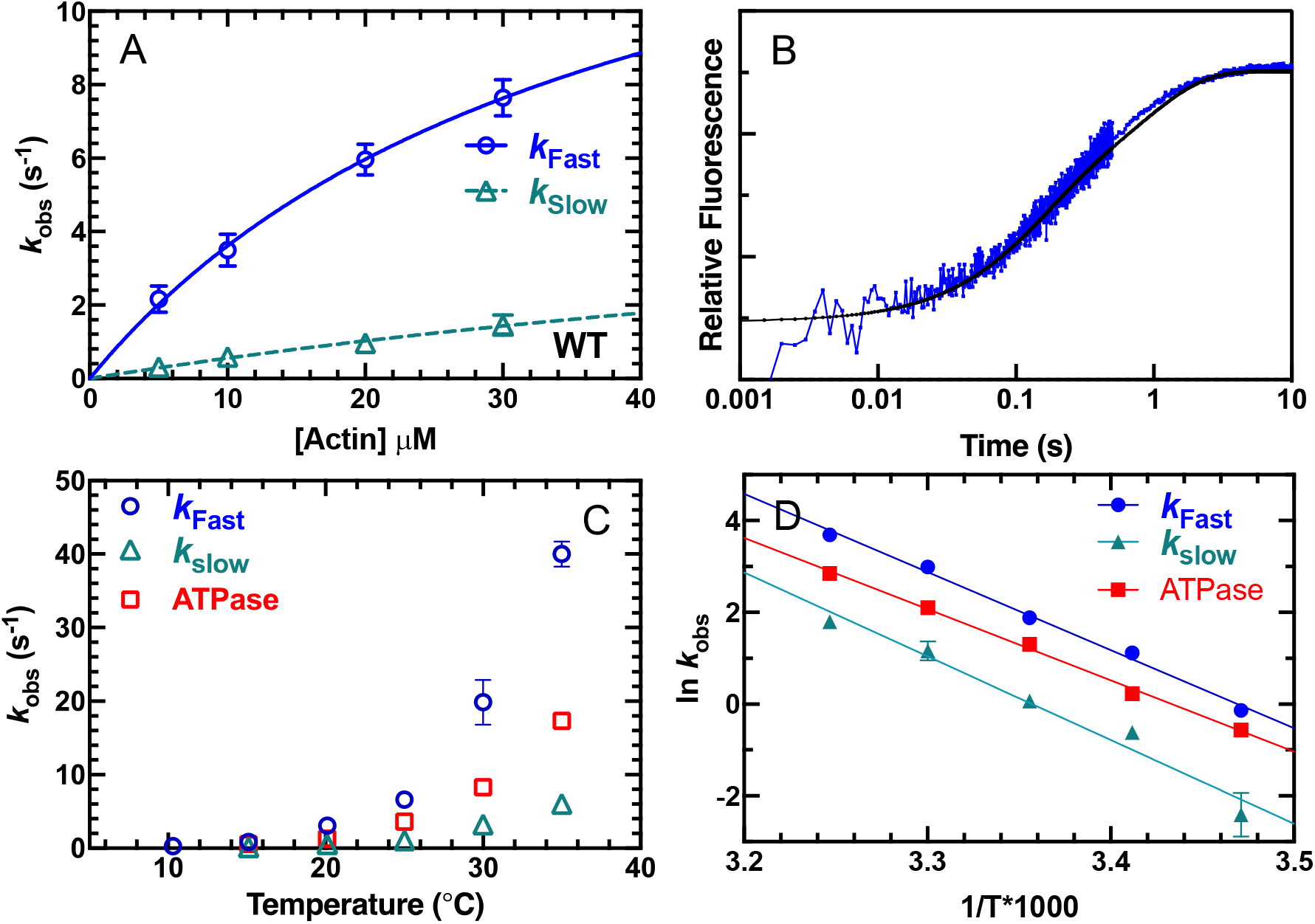
Measurements of actin-activation of the power stroke in M2β-S1. The rate constants for lever arm rotation during the power stroke were measured by monitoring the fluorescence enhancement of Alexa 488 during actin-activated product release. Sequential mix stopped-flow experiments were performed by mixing 0.25 µM M2β-S1 A488RLC with 0.2 µM Cy3ATP, aged for 10 s for hydrolysis to occur, and then mixed with varying concentrations of actin (5-30 µM). The fluorescence transients were best fit by a double exponential function. **(A)** The rate constants of the fast phase were plotted as a function of actin concentration and fit to a hyperbolic function. The slow phase was linearly dependent on actin-concentration and has a rate of 1.5 s^-1^ at 30 µM actin. Data points at each actin concentration represent the average ± SEM of 3-7 experiments from separate protein preparations. **(B)** Representative fluorescence transients in the presence of 30 µM actin (average of 2 transients) are shown fit to a double exponential function. **(C)** Experiments in panel A were performed as a function of temperature, and the rate constants of the fast and slow phases of the power stroke were plotted together with the actin-activated ATPase activity (10-35 °C). **(D)** Eyring plots of the power stroke rate constants and corresponding ATPase activity demonstrate the temperature dependence of the different rate constants (slopes, *k*_Fast_, -17.03 ± 0.76; *k*_Slow_, -18.25 ± 1.85; ATPase, -15.53 ± 0.51).

### Temperature dependence of phosphate release and power stroke

We performed sequential-mix experiments as described above to measure actin-activation of phosphate release and the power stroke under identical conditions, in the presence of 30 µM actin at three temperatures, 25, 30 and 35 °C (Figure 3 and Table S2). In phosphate release experiments, a lag was observed at 25 °C (8.2 ± 0.7 s^-1^) and 30 °C (21.6 ± 6.2 s^-1^), with an observed Pi release rate of 3.0 ± 0.1 s^-1^ and 7.1 ± 0.1 s^-1^, respectively. The fast phase of the power stroke was very similar to the lag in the Pi release measurements (7.3 ± 0.3 s^-1^ at 25 °C and 22.0 ± 1.4 s^-1^ at 30 °C), indicating that Pi release was rate-limited by the power stroke at these two temperatures. No lag was observed in the Pi release experiments at 35 °C, and the Pi release rate constant (19.2 ± 0.1 s^-1^) was nearly 50% slower than the fast power stroke rate constant (36.5 ± 1.3 s^-1^).

**Figure 3.**
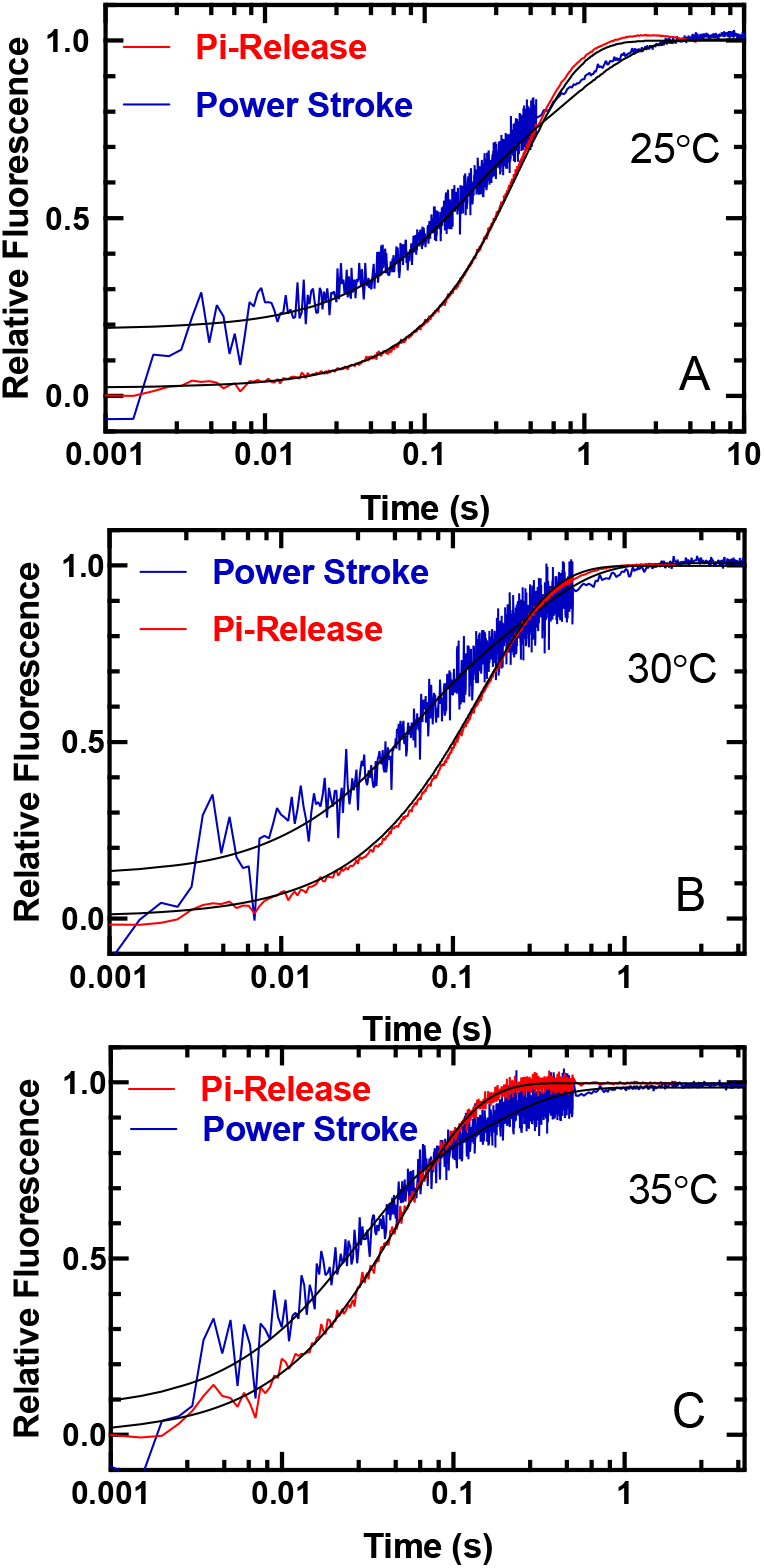
Temperature dependence of the power stroke and phosphate release in M2β-S1. The phosphate binding protein (MDCC-PBP) was used to monitor the phosphate release step using sequential mix experiments similar to that described in Figure 2. 1-2 µM M2β-S1 was mixed with substoichiometric ATP, aged for 10 s and then mixed with 30 µM actin and MDCC-PBP. Representative fluorescence transients (average of 2-3 normalized transients) of the power stroke (blue) and Pi release (red) in the presence of 30 µM actin are shown at **(A)** 25 °C and **(B)** 30 °C and **(C)** 35 °C. The power stroke experiments were best fit to a double exponential function at all temperatures. The Pi release experiments were best fit by a lag followed by a single exponential fluorescence increase at 25 and 30 °C, while the transients were single exponential at 35 °C. (25 °C, *k*_Fast_ = 7.3 ± 0.3 s^-1^, *k*_Slow_ = 1.3 ± 0.1 s^-1^, *k*_Lag_ = 8.2 ± 0.7 s^-1^, *k*_Pi_ = 3.0 ± 0.1 s^-1^, *A*_Fast_ = 0.5; 30 °C, *k*_Fast_ = 22 ± 1.4 s^-1^, *k*_Slow_ = 3.7 ± 0.2 s^-1^, *k*_Lag_ = 21.6 ± 6.2 s^-1^, *k*_Pi_ = 7.1 ± 0.1 s^-1^, *A*_Fast_ = 0.5; 35 °C, *k*_Fast_ = 36.5 ± 1.3 s^-1^, *k*_Slow_ = 5.0 ± 0.2 s^-1^, No Lag, *k*_Pi_ = 19.2 ± 0.1 s^-1^, *A*_Fast_ = 0.7).

### Impact of converter domain mutations on steady-state motor properties

We previously reported the steady-state and transient kinetic measurements of M2β-S1 F764L in our recent work (25). In the current work, we compared the steady-state motor properties of the two mutants with WT in parallel. Both R723G (6.1 ± 1.5 s^-1^) and F764L (6.4 ± 1.4 s^-1^) have slightly slower maximum ATPase rates compared with WT (8.3 ± 1.1 s^-1^), while the difference is not significant (Figure S5A and Table 2). The average sliding velocities in the *in vitro* motility assay were determined by pooling together 90 filaments from 3 protein preparations (30 each) at 0.24 µM loading concentration (Figure S5B). M2β-S1 R723G demonstrates a 7% increase in average sliding velocity (1699 ± 17 nm/s, *p* < 0.0001) compared with WT (1591 ± 16 nm/s), while F764L causes about a 10% decrease in average sliding velocity (1425 ± 16 nm/s, *p* < 0.0001) (Table 2).

### Transient kinetic measurements

We compared the transient kinetic parameters of M2β-S1 R723G and F764L mutants with WT (Table 2). The rate constants of WT and F764L were published in our previous work (25). We examined the ATP binding/hydrolysis steps in R723G by monitoring tryptophan fluorescence. An enhancement in fluorescence was observed upon mixing 1µM M2β-S1 R723G with varying concentrations of ATP (2.5-1000 µM) (Figure S6). The tryptophan fluorescence traces were best fit to a double exponential function with relative amplitude of the fast phase ∼60% of the signal at saturating ATP concentrations. The fast phase rate constants were hyperbolically dependent on ATP concentration, which allowed us to determine the maximum rate of ATP hydrolysis (231 ± 12 s^-1^) and ATP concentration dependence (*K*_0.5_ = 48 ± 9 µM) (Table 2). The second-order ATP binding constant in R723G determined by the linear dependence was 3.9 ± 0.1 µM^-1^·s^-1^ at low ATP concentrations. We also measured ATP-induced dissociation from actomyosin by mixing a M2β-S1:pyrene actin complex with varying concentrations of ATP (2 to 125 µM) (Figure S7). The pyrene fluorescence traces were best fit to a double exponential function. The rate constant of the fast phase (amplitude was ∼90% of the signal) plotted as a function of ATP concentration were fit to a hyperbolic equation *k*_obs_ = *K*’_1_*k*’_+2T_ × [ATP]/(1+*K*’_1_× [ATP])(25), which allowed us to determine the maximum rate *k*’_+2T_ = 668 ± 121 s^-1^ and second order binding constant for ATP *K*’_1_*k*’_+2T_ = 8.2 ± 0.1 µM (Table 2). The ADP release rate constant was measured by mixing an actomyosin.*mant*-ADP complex with 1mM ATP (data not shown). The fluorescence transients from three separate preparations were fit to a single exponential function, and the average and standard deviations were compared with WT and F764L (Table 2). The ADP release rate constant was also examined by mixing M2β-S1 R723G:pyrene actin.ADP complex (0.5 µM M2β-S1 R723G, 0.5 µM pyrene actin, and 50 µM ADP) with varying concentrations of ATP (10 to 2000 µM). The data were fit to a hyperbolic function and the maximum rate was defined as the ADP release rate constant as previously described (Table 2)(25). Overall, the rate constants measured for R723G were quite similar to WT except for the ADP release rate constant, which was slightly faster and consistent with increased velocity in the *in vitro* motility assay.

### Impact of mutations on structural kinetics of lever arm rotation

The steady-state FRET efficiency with Cy3ATP or Cy3ADP was determined for both R723G and F764L using M2β-S1 A488RLC constructs. No significant difference in FRET efficiency, distance, or ΔFRET (difference between Cy3ADP and Cy3ATP conditions) was found between R723G or F764L and WT (Figure 4 and Table 1). Measurements of the fast and slow power stroke rate constants showed that R723G causes a 37% decrease in the maximum rate of the fast phase (10.8 ± 4.1 s^-1^), while the decrease was slightly larger (50%) for F764L (8.7 ± 4.5 s^-1^) (Figure 5 and Table 2). Student’s *t*-tests were performed at each actin concentration to compare each mutant with WT, and significant differences were found at 30 µM actin in R723G, and 20 µM and 30 µM actin in F764L. The slow phase in the power stroke experiments was also actin-concentration dependent for R723G and F764L and was slower in both R723G (0.9 ± 0.2 s^-1^) and F764L (0.8 ± 0.3 s^-1^) compared with WT (1.5 ± 0.3 s^-1^) at 30 µM actin (Table 2). We estimated the duty ratio for the two mutant and WT constructs using an equation that assumes the hydrolysis and fast power stroke rate constants controls the transition into the strong binding states while the rate of ADP release from actomyosin controls the transition into the weak-binding states (see Methods). We found that compared to WT M2β-S1 (∼0.08) the estimated duty ratio is decreased in R723G (∼0.05) and F764L (∼0.06).

**Figure 4.**
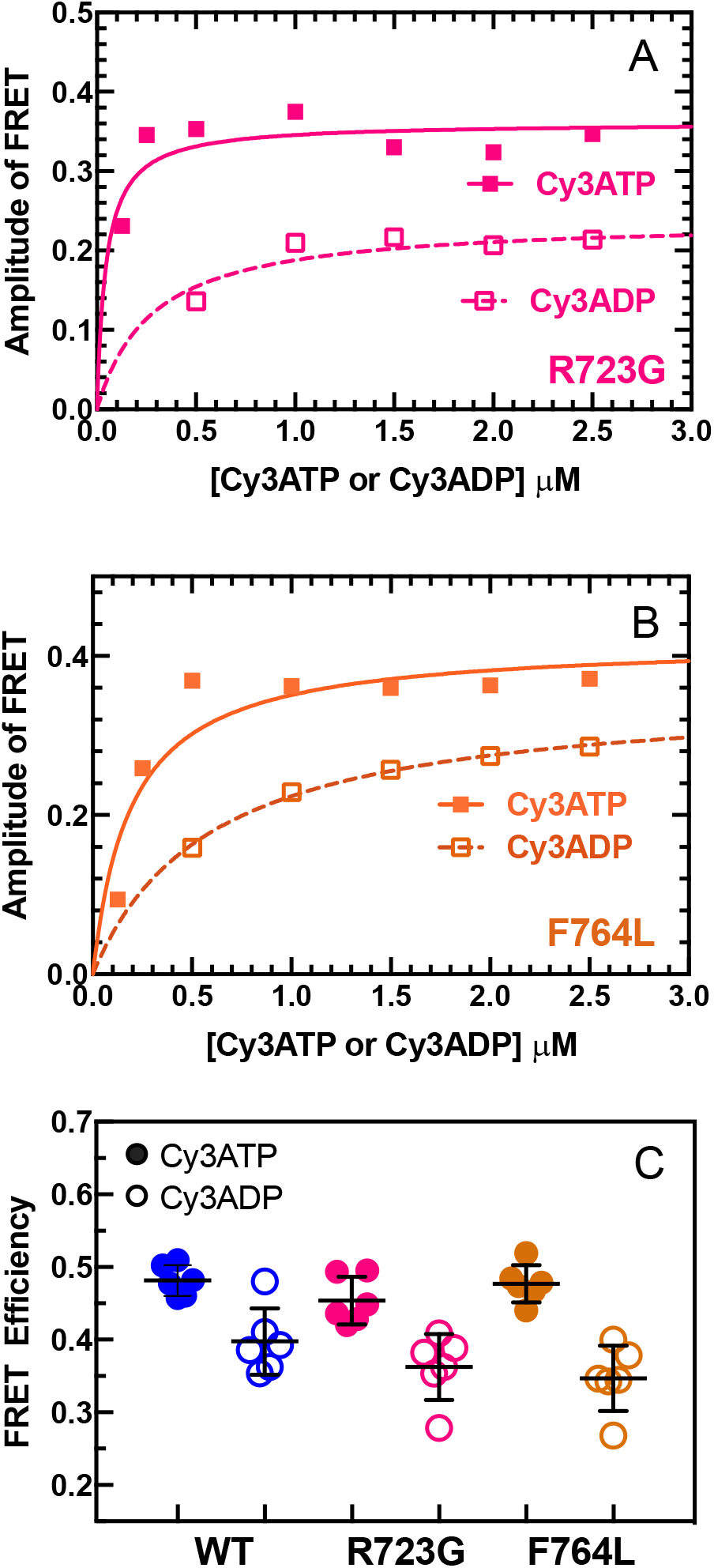
Impact of R723G and F764L mutants on lever arm conformation monitored by FRET. Measurements of Cy3ATP and Cy3ADP binding to M2β-S1 were examined as described in Figure 1. The amplitude of the FRET change was plotted as a function of Cy3ATP or Cy3ADP concentration and fit to a hyperbolic function for **(A)** R723G and **(B)** F764L. **(C)** Summary of steady-state FRET efficiency determined from measuring the emission spectra of 0.5 µM M2β-S1 in the presence of 1 µM Cy3ATP or Cy3ADP in WT, R723G, and F764L (average of 6 experiments from 3 protein preparations). The black bar represents the mean ± SD. Unpaired student’s *t*-tests were performed to compare each mutant with WT, and no significant differences were found between the mutants and WT.

**Figure 5.**
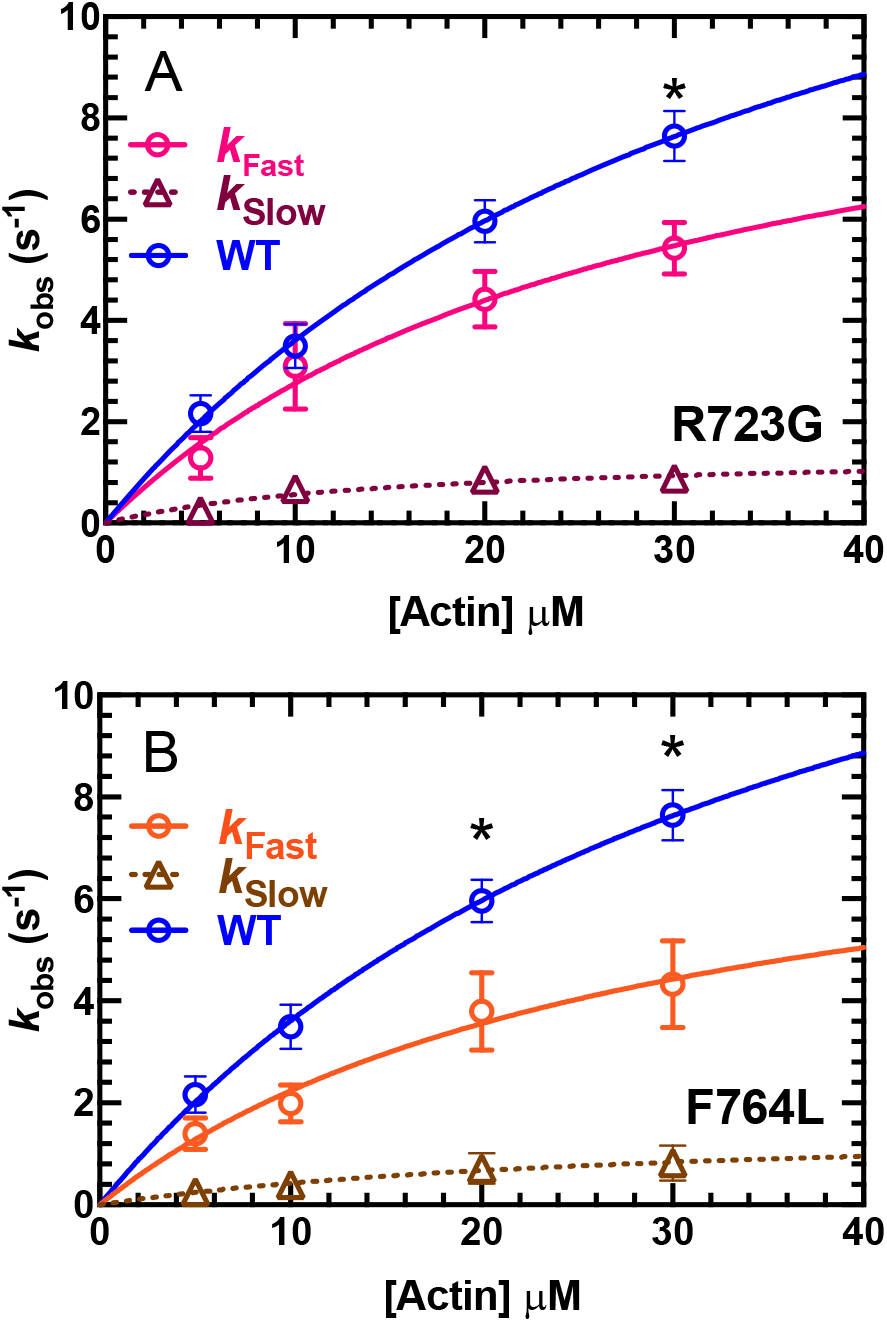
Impact of R723G and F764L mutants on the power stroke rate constant in M2β-S1. Sequential mix stopped-flow experiments were performed as described in Figure 2. The rate constants of the fast phase were plotted as a function of actin concentration and fit to a hyperbolic function to determine the maximum rate of power stroke for **(A)** R723G, 10.8 ± 4.1 s^-1^, or **(B)** F764L, 8.7 ± 4.5 s^-1^. The slow phase was also actin-concentration dependent for both mutants with a rate of 0.9 ± 0.2 s^-1^ for R723G, and 0.8 ± 0.3 s^-1^ for F764L at 30 µM actin. Data points at each actin concentration represent the average ± SEM of 3 experiments from separate protein preparations. The fast phase of the power stroke for WT is re-plotted from Figure 2 for comparison with the mutants. Student’s *t*-tests were performed at each actin concentration to compare each mutant with WT, and significant difference were found at 30 µM actin in R723G, and 20 µM and 30 µM actin in F764L (*, *p* < 0.05).

### Impact of converter domain mutations on mechanosensitivity

Additionally, we performed loaded *in vitro* motility assays to measure the sliding velocity in the presence of an increasing amount of tethering load induced by the presence of a motor-dead version of M2β-S1. We used a motor-dead mutant (E466A) M2β-S1, which is known to inhibit ATP hydrolysis in myosins (51) and movement of actin filaments in the motility assay. The advantage of using M2β-S1 E466A is that it has the same size and surface attachment strategy as all M2β-S1 constructs examined. The relative velocity (normalized to 0% motor-dead) was plotted as a function of the percentage of the motor-dead loaded on the motility surface (total concentration is constant at 0.6 µM). We observed a decrease in relative sliding velocity in both the WT and mutant M2β-S1 constructs with an increasing percentage of the motor-dead present. The relative reduction in sliding velocity was more pronounced and statistically different in R723G and F764L compared to WT at the highest motor-dead concentration (based on 3 separate protein preparations) (Figure 6).

**Figure 6.**
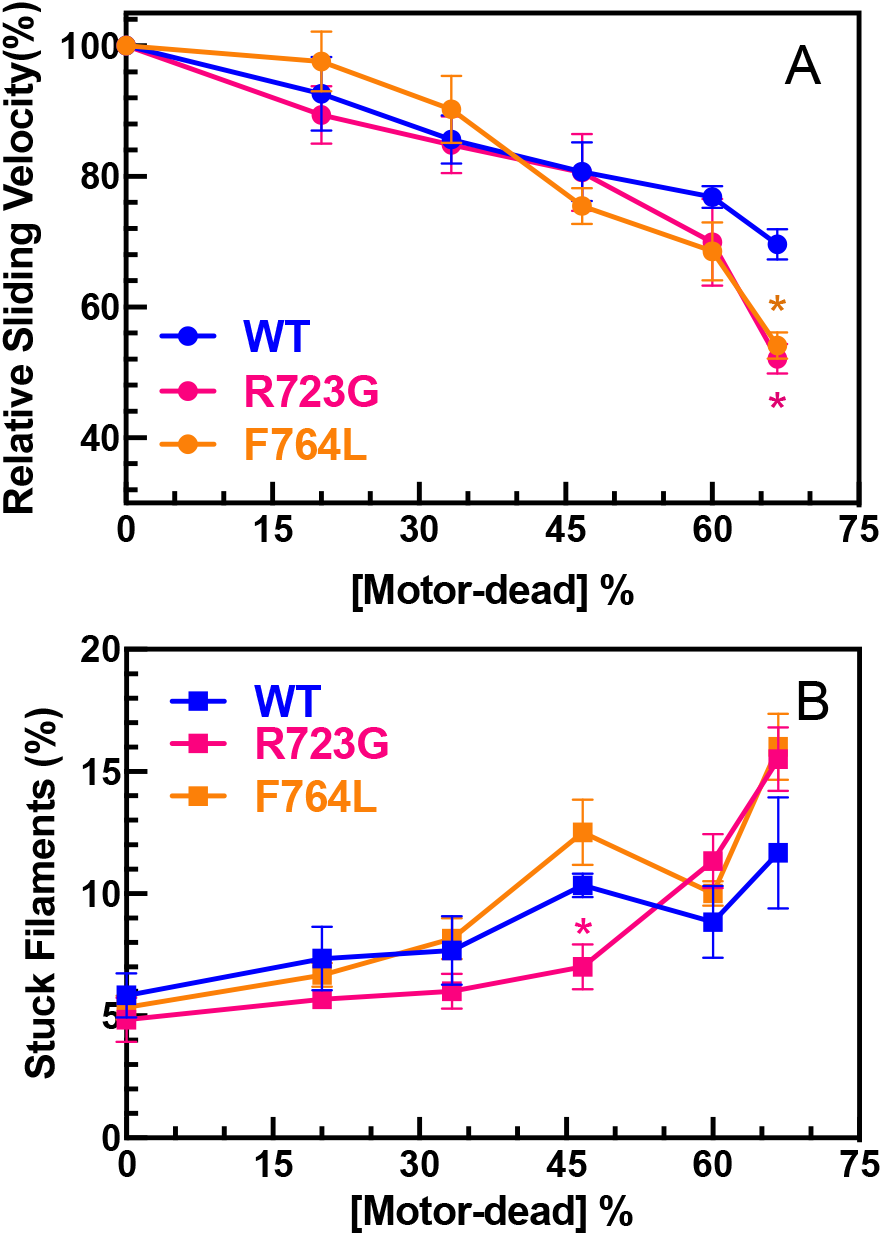
Loaded *in vitro* motility of M2β-S1 WT, R723G, and F764L. The impact of a tethering load imposed by the presence of M2β-S1 E466A (motor-dead), was examining in WT, R723G, and F764L M2β-S1 from 3 different protein preparations. The total concentration of M2β-S1 loaded onto the motility surface was kept constant at 0.6 µM. Velocities were analyzed by the program FAST(44). **(A)** The sliding velocity was expressed relative to 0% motor-dead and plotted as a function of the fraction of motor-dead present. **(B)** The percentage of filaments on the motility surface that were mobile was examined in each condition, and plotted as a function of the fraction motor-dead present. Each data point represents the mean ± SEM, *N* = 6. Unpaired student’s *t*-tests were performed to compare each mutant with WT at each [motor-dead]% (*, *p* < 0.05).

## Discussion

In the current work, we successfully utilized a FRET technique to monitor the structural kinetics of lever arm rotation (power stroke) in human cardiac myosin. We determined the power stroke rate constants in human cardiac myosin using sequential mix stopped-flow experiments, and the values agree well with the previous measurements in bovine cardiac myosin. We demonstrated clear evidence that the power stroke gates Pi release in human cardiac myosin, since the observed lag associated with Pi release was equivalent to the power stroke rate constant at lower temperatures (25 and 30 °C). At near physiological temperature (≥ 35°C), the power stroke is 2-fold faster than Pi release, and no lag was observed. We also measured the power stroke rate constants in two mutants, one associated with HCM and another associated with DCM, and measured their ensemble force generating capacity in a loaded *in vitro* motility assay. Surprisingly, both mutants slowed actin-activation of the power stroke and reduced ensemble force, indicating our measurements have revealed a critical mutation-specific defect that provides important insights into the molecular mechanism of inherited cardiomyopathies.

### The measurement of the power stroke in human cardiac myosin

In the power stroke measurements, two phases of the FRET transients were observed, a fast phase with a maximum rate constant of 17.2 ± 4.8 s^-1^, and a slow phase with a rate constant of 1.5 ± 0.3 s^-1^ at 30 µM actin. By plotting the fast and slow phase rate constants together with the ATPase rates in the presence of 30 µM actin at different temperatures (Figure 2B), we found the fast power stroke rate constant was faster than the ATPase rate at all temperatures examined, and the slow phase is slightly slower than the ATPase rate. Thus, we concluded that the fast phase rate constant represents the rate of the FRET change associated with the lever arm rotation (power stroke). The slow phase rate constant likely detected the FRET signal occurring between A488RLC and Cy3ADP when Cy3ADP was released from the nucleotide binding pocket.

The maximum rate of the power stroke in human cardiac myosin determined in the current study (17.2 s^-1^) agrees well with the power stroke measurement in bovine cardiac myosin (15.3 s^-1^) using a similar FRET technique (17). However, Woody et al. (11) recently reported a much faster working stroke rate, > 700 s^-1^, determined in an ultra-fast force clamp (UFFC)-equipped single-molecule optical trapping experiment. They monitored the actomyosin attachment duration as well as the detachment rate of human β-cardiac myosin heavy meromyosin (M2β HMM) under loaded conditions ranging from 1.5 pN to 4.5 pN, and the rate of the working stroke was found to be extremely fast (700 to 5,250 s^-1^) and increased with load.

It is well established that myosin, with the hydrolyzed products in its active site, associates weakly with actin, and then transitions into a strong actin-attached state to perform the power stroke. An initial actin-attached state has been proposed to be the pre-force generating intermediate before myosin transitions into the strong actomyosin state (11, 52, 53). The intermediate state is highly reversible (can rapidly detach from actin) but it is proposed to be stereospecific in nature (11). As Woody et al. (11) also suggested, the observed working stroke rate constant observed in their study represents an ensemble of several steps (actin-attachment followed by the power stroke and then Pi release), and also includes the rapid detachment from the intermediate state. The sequential mix stopped-flow experiments in our current work were done in the absence of load which may make it difficult to directly compare our power stroke rate constants to the optical trapping results. Indeed the detachment rate and power stroke reversal were found to be increased by load in the optical trapping experiments. In addition, only events that occur faster than the detachment rate were detected by the optical trap and included in the ensemble averaging.

### The sequence of events of the power stroke and Pi release in cardiac myosin

We observed a lag in the Pi release measurements that was similar to the power stroke rate constant measured by FRET at 25 °C and 30 °C. These results provide strong evidence that the power stroke occurs before Pi release in human cardiac myosin. To confirm this conclusion, we performed kinetic modeling using the ATPase pathway described in Scheme 1. Our kinetic simulation of the power stroke and Pi release fluorescence transients are best fit to a model with a power stroke rate constant of 25 s^-1^ that is followed by Pi release at 27.5 s^-1^ (Figure S8A&B and Table S3). Our proposed kinetic model (including most steps in the ATPase cycle) was also used to simulate the actin-activated ATPase data, and it matches well with our experimental results (Figure S8C and Table S3) (25).

Woody et al. (11) observed no effect on the rate of the ensemble average displacements with added 10 mM free Pi, also suggesting that Pi release occurs after the power stroke. Previous work found a two-step power stroke in myosin V, one faster step before Pi release and one slower step before ADP release (15). In both human and bovine cardiac myosins, only one-step was observed while the second step may be difficult to resolve since it may be similar to the nucleotide-release step, which we observe as our slower phase in the power stroke measurements. Nevertheless, the observed fast phase of the power stroke that occurs before Pi release in cardiac myosin is significantly slower than that observed in myosin V and skeletal muscle myosin (14, 15, 19). Cardiac muscle is known to have a slow rate of force development compared with skeletal muscle myosin at high Ca^2+^ concentrations, which suggests our power stroke measurements may correlate with the slower force development observed in cardiac muscle (54). The slower rate of force development may be crucial for tuning cooperative activation of the thin filament in cardiac muscle. Interestingly, activation of cardiac muscle is less dependent on Ca^2+^ concentrations compared with skeletal muscle (55). Force development may be an important regulatory step in cardiac muscle, which can be modified by additional Ca^2+^ independent thick-thin filament processes such as interactions with myosin binding protein C.

### Impact of mutations on the power stroke

Despite their clinical HCM and DCM phenotypes, both R723G and F764L mutations cause a significant decrease in the power stroke rate constant. Since in most models, the power stroke is tightly coupled to the weak-to-strong transition, we suggest that both mutations will slow the transition into the strong binding states. Structural analysis of the local environment of R723G revealed that it is a highly conserved arginine residue located at an extended loop in the converter domain, and is close to the ELC in the pre-power stroke state (Figure S9). The positive charge of the Arg-723 side chain forms a salt-bridge with the negatively charged side chain of glutamic acid at residue 136 in the ELC. This interaction may be crucial for the allosteric coupling between the converter domain and lever arm. Altering this interaction in myosin V impacts the ATP hydrolysis and recovery stroke rate constants (14, 56). The R723G mutation abolishes the charge-charge interaction between the converter domain and ELC, which may weaken the allosteric coupling between the two domains and destabilize the pre-power stroke conformation. The slower power stroke rate constant and faster ADP release rate constant determined in the current work lead to a decreased duty ratio in R723G. Indeed, in a recent study comparing several HCM mutations, R723G is associated with a lower duty ratio under both unloaded and loaded conditions (35). Our results of a slower power stroke and faster ADP release, which shorten actomyosin attachment duration, agree with these conclusions.

As described in our recent work (14), residue Phe-764 is located at a hydrophobic patch between the relay helix and the converter domain. The conversion of leucine to phenylalanine at residue 764 likely weakens the hydrophobic interaction and increases the flexibility of the local environment, which may decrease the communication efficiency between the converter domain and the active site. The slowing of the power stroke rate constant in F764L also contributes to a decreased duty ratio while the slower ADP release rate constant partially offsets the impact of the slower power stroke. Overall, the F764L mutant likely disrupts communication between subdomains within the motor, which may contribute to its impairment of the structural kinetics of lever arm rotation during the power stroke.

Additionally, we observed no significant difference in the steady-FRET efficiency in either R723G or F764L, indicating similar pre- and post-power stroke conformations of the lever arm in both R723G and F764L in the absence of actin (Table 1). It would be interesting to investigate further whether the two mutations shift the mole fraction of pre- and post-power stroke structural states in the presence and absence of actin, which can be determined with time-resolved fluorescence energy transfer (TR-FRET) measurements (14).

### Impact of mutations on load sensitivity

External load is thought to alter the ATPase kinetics of all myosins (57). Cardiac myosin, working as an ensemble to generate force, can alter its force-velocity (*F*-*v*) properties (power output) under various loaded conditions. The presence of high loads inhibits the release of ADP, the load sensing step during the ATPase cycle, to prolong the actomyosin attachment duration and slow down sliding velocity (58). The loaded *in vitro* motility assay is designed to examine the impact of physiological loads, as the myosin molecules on the motility surface have to overcome the tethering force associated with an actin binding protein or motor-dead myosin (59, 60).

The converter domain is the load-sensing region of the myosin motor that is thought to communicate external load to the nucleotide binding pocket, which alters ADP release kinetics. Variations in the converter domain of myosin have been shown to modulate the contractile properties of various muscle types (61). Interestingly, both converter domain mutations R723G and F764L showed a trend of a reduced capability to overcome the tethering load compared with WT, and the trend is similar in the F764L and R723G mutants. In contrast, the unloaded sliding velocity is 7% faster in R723G, and 10% slower in F764L compared with WT, respectively. In our previous study that examined the two corresponding mutations (R712G and F750L) in myosin V, an even larger difference was observed in the ability of the mutations to overcome resistive load (e.g., assessed by the number of filaments moving as a function of motor-dead% on the surface).

Notably, R723G and F764L mutations both alter the ADP release rate constant determined with transient kinetic studies, which correlates with the changes observed in unloaded *in vitro* motility (Table 2). The faster ADP release rate constant (218±15 s^-1^) observed in R723G correlates well with its faster unloaded sliding velocity, and the slower ADP release rate constant in F764L (132±16 s^-1^) also agrees with its slower sliding velocity compared with WT (174±12 s^-1^). But given the trend of reduced ability of the two mutants to overcome external load, we hypothesize that under loaded conditions, the changes in the kinetics of individual steps of the ATPase cycle could be even more significant. In a muscle force-velocity relationship, R723G would be expected to increase unloaded shortening velocity but reduce isometric force, while the F764L mutant would be expected to reduce unloaded sliding velocity and isometric force. The reduced duty ratio in both R723G and F764L may contribute to their reduced ensemble force production. Interestingly, R723G was found to cause a reduction in intrinsic force measured with single molecule optical trapping studies (34, 35), and this could also contribute to the observed reduction in ensemble force. The intrinsic force in F764L has not been measured, but its more severely reduced power stroke rate constant compared to R723G suggests this parameter is an important determinant of duty ratio and ensemble force production.

### Potential disease-causing mechanisms for R723G and F764L

The R723G mutation causes a slightly lower maximum ATPase, reduced power stroke rate constant, reduced intrinsic force, lower duty ratio, and reduced ensemble force production. All the changes together indicate that R723G impairs motor contractility, which contrasts with the proposed hypercontractility hypothesis of HCM mutations (62). The F764L mutation has a reduced maximum ATPase, sliding velocity, duty ratio, and power stroke rate constant which agrees with the hypocontractility hypothesis for DCM mutations. However, the changes in many of these parameters are relatively small.

A recent ‘mesa’ theory proposes that residues located on a plateau-like surface within the cardiac myosin motor domain are important for forming an auto-inhibited structure called the interacting-heads motif (IHM)(63-65). Myosin heads in the IHM contain head-head and head-tail interactions that are proposed to cause a 10-fold slower ATP turnover rate and inhibit interactions with actin until they transition out of the folded OFF state. Mutations located on the mesa surface, including R723G, may destabilize the IHM structure and increase the number of active heads on thick filaments (34, 63). Therefore, the R723G mutation may increase the total number of available myosin heads on the thick filaments and thus increase overall isometric force produced in muscle, despite its reduced intrinsic motor properties. Phe-764 is not predicted to be at the interface mediating the formation of the IHM, but may still indirectly promote the SRX by stabilizing a conformation required for entry into the IHM.

## Conclusion

In conclusion, cardiomyopathy mutants that are associated with HCM and DCM phenotypes can directly alter the power stroke rate constant, a key step in the force generating ATPase cycle of myosins. Future single molecule optical trapping experiments to characterize the impact of these mutants under loaded conditions could shed light on the importance of load-sensitivity in HCM and DCM. In addition, examining the impact of these two converter domain mutations on the stability of the SRX state with M2β HMM constructs, using single turnover and electron microscopy experiments, will be extremely revealing. Overall, our results provide evidence that examining the impact of mutations on important conformational changes can uncover specific structural defects and may lead novel insight into disease mechanisms and therapeutic interventions.

---

**Scheme 1.**
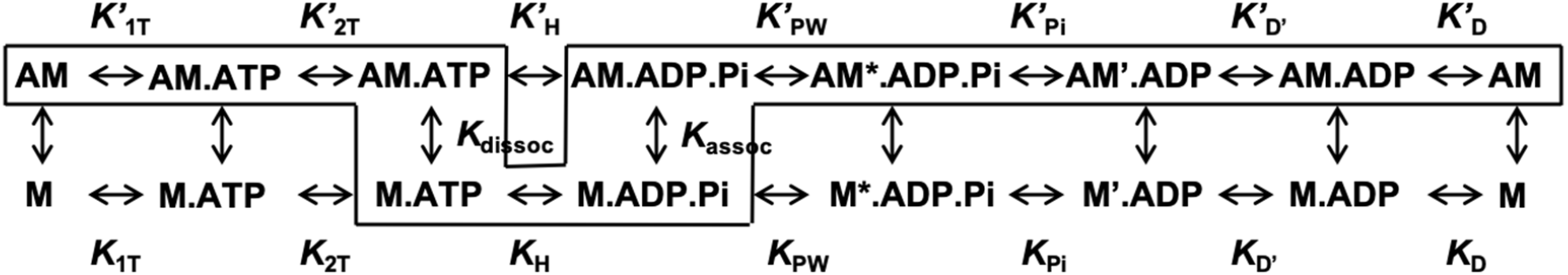
Actomyosin ATPase pathway.

## Author contributions

Wanjian Tang and Christopher Yengo designed and performed experiments. Jinghua Ge helped with the expression of human V105C regulatory light chain. William C. Unrath and Rohini Desetty provided support in molecular cloning, virus amplification, and cell culture. The text was co-written by Wanjian Tang and Christopher M. Yengo.

## Acknowledgements

This works was supported by a NIH grant to C.M.Y. (HL127699) and an AHA grant to W.T (19PRE34380569).

## References

1. Geeves, M. A. 2016. Review: The ATPase mechanism of myosin and actomyosin: The ATPase Mechanism of Myosin and Actomyosin. Biopolymers 105(8):483–491.

2. Málnási-Csizmadia, A., and M. Kovács. 2010. Emerging complex pathways of the actomyosin powerstroke. Trends in Biochemical Sciences 35(12):684–690.

3. Sweeney, H. L., and A. Houdusse. 2010. Structural and functional insights into the Myosin motor mechanism. Annu Rev Biophys 39:539–557.

4. Preller, M., and D. J. Manstein. 2013. Myosin structure, allostery, and mechano-chemistry. Structure 21(11):1911–1922.

5. Llinas, P., T. Isabet, L. Song, V. Ropars, B. Zong, H. Benisty, S. Sirigu, C. Morris, C. Kikuti, D. Safer, H. L. Sweeney, and A. Houdusse. 2015. How actin initiates the motor activity of Myosin. Dev Cell 33(4):401–412.

6. Lymn, R. W., and E. W. Taylor. 1971. Mechanism of adenosine triphosphate hydrolysis by actomyosin. Biochemistry 10(25):4617–4624.

7. Veigel, C., and C. F. Schmidt. 2011. Moving into the cell: single-molecule studies of molecular motors in complex environments. Nat Rev Mol Cell Biol 12(3):163–176.

8. Kodera, N., and T. Ando. 2014. The path to visualization of walking myosin V by high-speed atomic force microscopy. Biophys Rev 6(3-4):237–260.

9. Mentes, A., A. Huehn, X. Liu, A. Zwolak, R. Dominguez, H. Shuman, E. M. Ostap, and C. V. Sindelar. 2018. High-resolution cryo-EM structures of actin-bound myosin states reveal the mechanism of myosin force sensing. Proc Natl Acad Sci U S A 115(6):1292–1297.

10. Shuman, H., M. J. Greenberg, A. Zwolak, T. Lin, C. V. Sindelar, R. Dominguez, and E. M. Ostap. 2014. A vertebrate myosin-I structure reveals unique insights into myosin mechanochemical tuning. Proc Natl Acad Sci U S A 111(6):2116–2121.

11. Woody, M. S., D. A. Winkelmann, M. Capitanio, E. M. Ostap, and Y. E. Goldman. 2019. Single molecule mechanics resolves the earliest events in force generation by cardiac myosin. Elife 8.

12. Wulf, S. F., V. Ropars, S. Fujita-Becker, M. Oster, G. Hofhaus, L. G. Trabuco, O. Pylypenko, H. L. Sweeney, A. M. Houdusse, and R. R. Schroder. 2016. Force-producing ADP state of myosin bound to actin. Proc Natl Acad Sci U S A 113(13):E1844–1852.

13. Greenberg, M. J., T. Lin, H. Shuman, and E. M. Ostap. 2015. Mechanochemical tuning of myosin-I by the N-terminal region. Proc Natl Acad Sci U S A 112(26):E3337–3344.

14. Gunther, L. K., J. A. Rohde, W. Tang, S. D. Walton, W. C. Unrath, D. V. Trivedi, J. M. Muretta, D. D. Thomas, and C. M. Yengo. 2019. Converter domain mutations in myosin alter structural kinetics and motor function. Journal of Biological Chemistry 294(5):1554–1567.

15. Trivedi, D. V., J. M. Muretta, A. M. Swenson, J. P. Davis, D. D. Thomas, and C. M. Yengo. 2015. Direct measurements of the coordination of lever arm swing and the catalytic cycle in myosin V. Proc Natl Acad Sci U S A 112(47):14593–14598.

16. Rohde, J. A., O. Roopnarine, D. D. Thomas, and J. M. Muretta. 2018. Mavacamten stabilizes an autoinhibited state of two-headed cardiac myosin. Proc Natl Acad Sci U S A 115(32):E7486–E7494.

17. Rohde, J. A., D. D. Thomas, and J. M. Muretta. 2017. Heart failure drug changes the mechanoenzymology of the cardiac myosin powerstroke. Proc Natl Acad Sci U S A 114(10):E1796–E1804.

18. Muretta, J. M., K. J. Petersen, and D. D. Thomas. 2013. Direct real-time detection of the actin-activated power stroke within the myosin catalytic domain. Proc Natl Acad Sci U S A 110(18):7211–7216.

19. Muretta, J. M., J. A. Rohde, D. O. Johnsrud, S. Cornea, and D. D. Thomas. 2015. Direct real-time detection of the structural and biochemical events in the myosin power stroke. Proc Natl Acad Sci U S A 112(46):14272–14277.

20. Moore, J. R., L. Leinwand, and D. M. Warshaw. 2012. Understanding cardiomyopathy phenotypes based on the functional impact of mutations in the myosin motor. Circulation Research 111(3):375–385.

21. Enjuto, M., A. Francino, F. Navarro-Lopez, D. Viles, J. C. Pare, and A. M. Ballesta. 2000. Malignant hypertrophic cardiomyopathy caused by the Arg723Gly mutation in beta-myosin heavy chain gene. J Mol Cell Cardiol 32(12):2307–2313.

22. Kamisago, M., S. D. Sharma, S. R. DePalma, S. Solomon, P. Sharma, B. McDonough, L. Smoot, M. P. Mullen, P. K. Woolf, E. D. Wigle, J. G. Seidman, and C. E. Seidman. 2000. Mutations in sarcomere protein genes as a cause of dilated cardiomyopathy. N Engl J Med 343(23):1688–1696.

23. Baumketner, A. 2012. The mechanism of the converter domain rotation in the recovery stroke of myosin motor protein. Proteins: Structure, Function and Bioinformatics 80(12):2701–2710.

24. Bloemink, M. J., G. C. Melkani, S. I. Bernstein, and M. A. Geeves. 2016. The relay/converter interface influences hydrolysis of ATP by skeletal muscle myosin II. Journal of Biological Chemistry 291(4):1763–1773.

25. Tang, W., W. C. Unrath, R. Desetty, and C. M. Yengo. 2019. Dilated cardiomyopathy mutation in the converter domain of human cardiac myosin alters motor activity and response to omecamtiv mecarbil. J Biol Chem 294(46):17314–17325.

26. Ujfalusi, Z., C. D. Vera, S. M. Mijailovich, M. Svicevic, E. C. Yu, M. Kawana, K. M. Ruppel, J. A. Spudich, M. A. Geeves, and L. A. Leinwand. 2018. Dilated cardiomyopathy myosin mutants have reduced force-generating capacity. Journal of Biological Chemistry 293(23):9017–9029.

27. Debold, E. P., J. P. Schmitt, J. B. Patlak, S. E. Beck, J. R. Moore, J. G. Seidman, C. Seidman, and D. M. Warshaw. 2007. Hypertrophic and dilated cardiomyopathy mutations differentially affect the molecular force generation of mouse alpha-cardiac myosin in the laser trap assay. Am J Physiol Heart Circ Physiol 293(1):H284–291.

28. Schmitt, J. P., E. P. Debold, F. Ahmad, A. Armstrong, A. Frederico, D. A. Conner, U. Mende, M. J. Lohse, D. Warshaw, C. E. Seidman, and J. G. Seidman. 2006. Cardiac myosin missense mutations cause dilated cardiomyopathy in mouse models and depress molecular motor function. Proceedings of the National Academy of Sciences 103(39):14525–14530.

29. Palmer, B. M., J. P. Schmitt, C. E. Seidman, J. G. Seidman, Y. Wang, S. P. Bell, M. M. LeWinter, and D. W. Maughan. 2013. Elevated rates of force development and MgATP binding in F764L and S532P myosin mutations causing dilated cardiomyopathy. Journal of Molecular and Cellular Cardiology 57(1):23–31.

30. Kirschner, S. E., E. Becker, M. Antognozzi, H. P. Kubis, A. Francino, F. Navarro-Lopez, N. Bit-Avragim, A. Perrot, M. M. Mirrakhimov, K. J. Osterziel, W. J. McKenna, B. Brenner, and T. Kraft. 2005. Hypertrophic cardiomyopathy-related beta-myosin mutations cause highly variable calcium sensitivity with functional imbalances among individual muscle cells. Am J Physiol Heart Circ Physiol 288(3):H1242–1251.

31. Kohler, J., G. Winkler, I. Schulte, T. Scholz, W. McKenna, B. Brenner, and T. Kraft. 2002. Mutation of the myosin converter domain alters cross-bridge elasticity. Proc Natl Acad Sci U S A 99(6):3557–3562.

32. Seebohm, B., F. Matinmehr, J. Kohler, A. Francino, F. Navarro-Lopez, A. Perrot, C. Ozcelik, W. J. McKenna, B. Brenner, and T. Kraft. 2009. Cardiomyopathy mutations reveal variable region of myosin converter as major element of cross-bridge compliance. Biophys J 97(3):806–824.

33. Kraft, T., E. R. Witjas-Paalberends, N. M. Boontje, S. Tripathi, A. Brandis, J. Montag, J. L. Hodgkinson, A. Francino, F. Navarro-Lopez, B. Brenner, G. J. Stienen, and J. van der Velden. 2013. Familial hypertrophic cardiomyopathy: functional effects of myosin mutation R723G in cardiomyocytes. J Mol Cell Cardiol 57:13–22.

34. Kawana, M., S. S. Sarkar, S. Sutton, K. M. Ruppel, and J. A. Spudich. 2017. Biophysical properties of human beta-cardiac myosin with converter mutations that cause hypertrophic cardiomyopathy. Sci Adv 3(2):e1601959.

35. Vera, C. D., C. A. Johnson, J. Walklate, A. Adhikari, M. Svicevic, S. M. Mijailovich, A. C. Combs, S. J. Langer, K. M. Ruppel, J. A. Spudich, M. A. Geeves, and L. A. Leinwand. 2019. Myosin motor domains carrying mutations implicated in early or late onset hypertrophic cardiomyopathy have similar properties. J Biol Chem 294(46):17451–17462.

36. White, H. D., B. Belknap, and M. R. Webb. 1997. Kinetics of nucleoside triphosphate cleavage and phosphate release steps by associated rabbit skeletal actomyosin, measured using a novel fluorescent probe for phosphate. Biochemistry 36(39):11828–11836.

37. Winkelmann, D. A., E. Forgacs, M. T. Miller, and A. M. Stock. 2015. Structural basis for drug-induced allosteric changes to human beta-cardiac myosin motor activity. Nat Commun 6:7974.

38. Nag, S., D. V. Trivedi, S. S. Sarkar, A. S. Adhikari, M. S. Sunitha, S. Sutton, K. M. Ruppel, and J. A. Spudich. 2017. The myosin mesa and the basis of hypercontractility caused by hypertrophic cardiomyopathy mutations. Nat Struct Mol Biol 24(6):525–533.

39. Pardee, J. D., and J. A. Spudich. 1982. Purification of muscle actin. Methods Enzymol 85 Pt B:64–181.

40. Pollard, T. D. 1984. Polymerization of ADP-actin. J Cell Biol 99(3):769–777.

41. Swenson, A. M., W. Tang, C. A. Blair, C. M. Fetrow, W. C. Unrath, M. J. Previs, K. S. Campbell, and C. M. Yengo. 2017. Omecamtiv Mecarbil Enhances the Duty Ratio of Human beta-Cardiac Myosin Resulting in Increased Calcium Sensitivity and Slowed Force Development in Cardiac Muscle. J Biol Chem 292(9):3768–3778.

42. De La Cruz, E. M., H. L. Sweeney, and E. M. Ostap. 2000. ADP Inhibition of Myosin V ATPase Activity. Biophysical Journal 79(3):1524–1529.

43. Meijering, E., O. Dzyubachyk, and I. Smal. 2012. Methods for cell and particle tracking. Methods Enzymol 504:183–200.

44. Aksel, T., E. Choe Yu, S. Sutton, K. M. Ruppel, and J. A. Spudich. 2015. Ensemble force changes that result from human cardiac myosin mutations and a small-molecule effector. Cell Rep 11(6):910–920.

45. Jacobs, D. J., D. Trivedi, C. David, and C. M. Yengo. 2011. Kinetics and thermodynamics of the rate-limiting conformational change in the actomyosin V mechanochemical cycle. J Mol Biol 407(5):716–730.

46. Lakowicz, J. R. 2006. Principles of Fluorescence Spectroscopy. Springer, Boston, MA.

47. Johnson, K. A., Z. B. Simpson, and T. Blom. 2009. Global Kinetic Explorer: A new computer program for dynamic simulation and fitting of kinetic data. Analytical Biochemistry 387(1):20–29.

48. Johnson, K. A., Z. B. Simpson, and T. Blom. 2009. FitSpace explorer: an algorithm to evaluate multidimensional parameter space in fitting kinetic data. Anal Biochem 387(1):30–41.

49. Forgacs, E., T. Sakamoto, S. Cartwright, B. Belknap, M. Kovacs, J. Toth, M. R. Webb, J. R. Sellers, and H. D. White. 2009. Switch 1 mutation S217A converts myosin V into a low duty ratio motor. J Biol Chem 284(4):2138–2149.

50. Liu, Y., H. D. White, B. Belknap, D. A. Winkelmann, and E. Forgacs. 2015. Omecamtiv Mecarbil Modulates the Kinetic and Motile Properties of Porcine β-Cardiac Myosin. Biochemistry 54(10):1963–1975.

51. Yengo, C. M., E. M. De la Cruz, D. Safer, E. M. Ostap, and H. L. Sweeney. 2002. Kinetic characterization of the weak binding states of myosin V. Biochemistry 41(26):8508–8517.

52. Gunther, L. K., J. A. Rohde, W. Tang, J. A. Cirilo, Jr., C. P. Marang, B. D. Scott, D. D. Thomas, E. P. Debold, and C. M. Yengo. 2020. FRET and optical trapping reveal mechanisms of actin-activation of the power stroke and phosphate-release in myosin V. J Biol Chem.

53. Sun, M., M. B. Rose, S. K. Ananthanarayanan, D. J. Jacobs, and C. M. Yengo. 2008. Characterization of the pre-force-generation state in the actomyosin cross-bridge cycle. Proc Natl Acad Sci U S A 105(25):8631–8636.

54. Regnier, M., H. Martin, R. J. Barsotti, A. J. Rivera, D. A. Martyn, and E. Clemmens. 2004. Cross-bridge versus thin filament contributions to the level and rate of force development in cardiac muscle. Biophys J 87(3):1815–1824.

55. Hancock, W. O., L. L. Huntsman, and A. M. Gordon. 1997. Models of calcium activation account for differences between skeletal and cardiac force redevelopment kinetics. J Muscle Res Cell Motil 18(6):671–681.

56. De La Cruz, E. M., A. L. Wells, H. L. Sweeney, and E. M. Ostap. 2000. Actin and light chain isoform dependence of myosin V kinetics. Biochemistry 39(46):14196–14202.

57. Greenberg, M. J., G. Arpag, E. Tuzel, and E. M. Ostap. 2016. A Perspective on the Role of Myosins as Mechanosensors. Biophys J 110(12):2568–2576.

58. Greenberg, M. J., H. Shuman, and E. M. Ostap. 2014. Inherent force-dependent properties of beta-cardiac myosin contribute to the force-velocity relationship of cardiac muscle. Biophys J 107(12):L41–L44.

59. Greenberg, M. J., and J. R. Moore. 2010. The molecular basis of frictional loads in the in vitro motility assay with applications to the study of the loaded mechanochemistry of molecular motors. Cytoskeleton (Hoboken) 67(5):273–285.

60. Nag, S., R. F. Sommese, Z. Ujfalusi, A. Combs, S. Langer, S. Sutton, L. A. Leinwand, M. A. Geeves, K. M. Ruppel, and J. A. Spudich. 2015. Contractility parameters of human beta-cardiac myosin with the hypertrophic cardiomyopathy mutation R403Q show loss of motor function. Sci Adv 1(9):e1500511.

61. Swank, D. M., A. F. Knowles, J. A. Suggs, F. Sarsoza, A. Lee, D. W. Maughan, and S. I. Bernstein. 2002. The myosin converter domain modulates muscle performance. Nat Cell Biol 4(4):312–316.

62. Spudich, J. A. 2014. Hypertrophic and dilated cardiomyopathy: Four decades of basic research on muscle lead to potential therapeutic approaches to these devastating genetic diseases. Biophysical Journal 106(6):1236–1249.

63. Alamo, L., J. S. Ware, A. Pinto, R. E. Gillilan, J. G. Seidman, C. E. Seidman, and R. Padron. 2017. Effects of myosin variants on interacting-heads motif explain distinct hypertrophic and dilated cardiomyopathy phenotypes. Elife 6.

64. Hooijman, P., M. A. Stewart, and R. Cooke. 2011. A new state of cardiac myosin with very slow ATP turnover: a potential cardioprotective mechanism in the heart. Biophys J 100(8):1969–1976.

65. Spudich, J. A. 2015. The myosin mesa and a possible unifying hypothesis for the molecular basis of human hypertrophic cardiomyopathy. Biochem. Soc. Trans. 43(1):64–72.

