## Supplemental Information for "Cardiomyopathy Mutations Impact the Power Stroke of Human Cardiac Myosin"

Table **S1**. ATPase and motility comparison of M2β-S1 A488RLC, Non-Ex and hRLC-unlabeled.

| **Steady-State ATPase Values, (Mean ± SEM)** | **M2β-S1 Non-Ex** $N = 3$ | **M2β-S1 hRLC** $N = 1$ | **M2β-S1 A488 RLC** $N = 3$ |
|---|---|---|---|
| $v_0$ ($s^{-1}$) | 0.02 ± 0.01 | 0.03 | 0.01 ± 0.01 |
| $k_{cat}$ ($s^{-1}$) | 5.4 ± 1.3 | 5.3 ± 0.9 | 4.4 ± 0.6 |
| $K_{ATPase}$ (μM) | 53 ± 22 | 31 ± 11 | 32 ± 9 |
| ***In vitro* motility (Mean ± SEM)** $N = 50$ | | | |
| Mean Velocity (nm/s) | 1520 ± 25 | 1300 ± 14 | 1394 ± 29 |

Table **S2**. Temperature dependence of Pi release and power stroke.

| | Power stroke ($s^{-1}$) | | Pi release ($s^{-1}$) | |
|---|---|---|---|---|
| Temperature (°C) | $\boldsymbol{k}_{\mathbf{Fast}}$ | $\boldsymbol{k}_{\mathbf{Slow}}$ | $\boldsymbol{k}_{\mathbf{Lag}}$ | $\boldsymbol{k}_{\mathbf{Pi}}$ |
| 25 | 7.3 ± 0.3 | 1.3 ± 0.1 | 8.2 ± 0.7 | 3.0 ± 0.1 |
| 30 | 22 ± 1.4 | 3.7 ± 0.2 | 21.6 ± 6.2 | 7.1 ± 0.1 |
| 35 | 36.5 ± 1.3 | 5.0 ± 0.2 | No lag | 19.2 ± 0.1 |

Table **S3**. Rate constants used for power stroke, Pi release, and ATPase simulations.

| Rate constants | WT | |
|---|---|---|
| | Forward | Reverse |
| $k'_{1T}$ | 20 μ$M^{-1}\cdot s^{-1}$ | 1000 $s^{-1}$ |
| $k'_{2T}$ ($s^{-1}$) | 512 | 0.001 |
| $k_{dissoc}$ ($s^{-1}$) | 1000 | 10 |
| $k_H$ ($s^{-1}$) | 120 | 60 |
| $k_{assoc}$ | 5-7.5 μ$M^{-1}\cdot s^{-1}$ | 1000 $s^{-1}$ |
| $k_{PWF}$ ($s^{-1}$) | 25 | 1 |
| $k_{Pi}$ ($s^{-1}$) | 27.5 | 0.1 |
| $k'_{D'}$ ($s^{-1}$) | 500 | 1 |
| $k'_D$ ($s^{-1}$) | 180 | 1 |
| $k_{cat}$ ($s^{-1}$) | 7.2 ± 0.3 | |
| $K_{ATPase}$ (μM) | 59.8 ± 3.5 | |

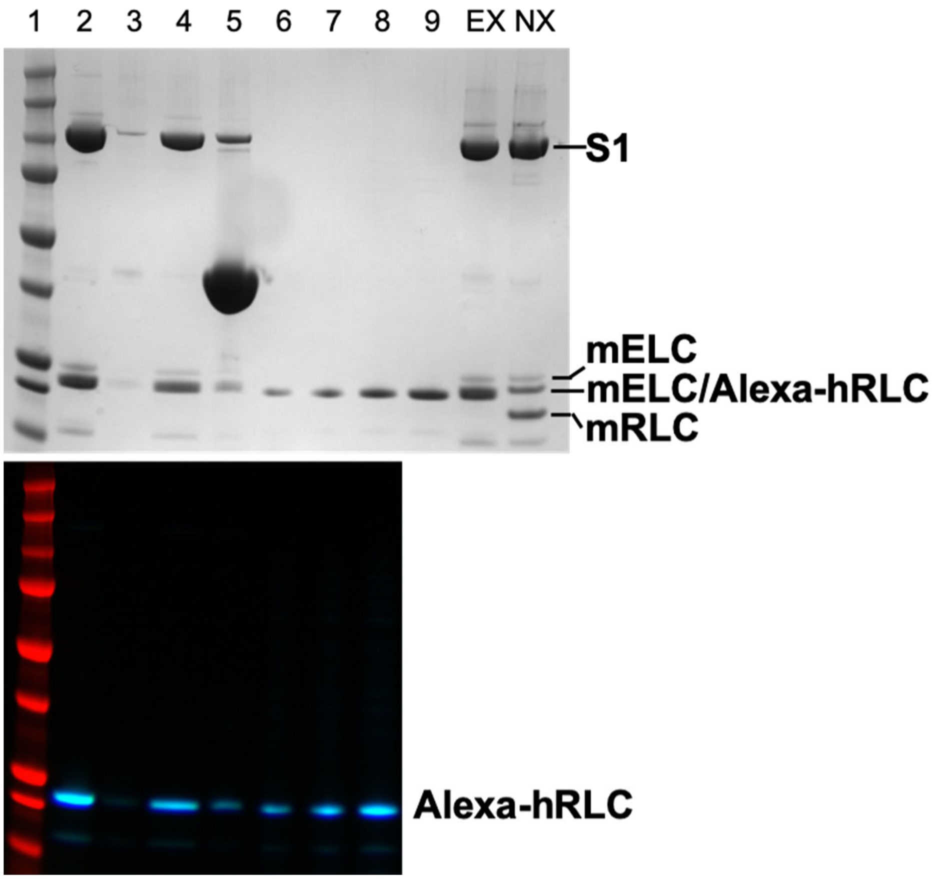


Figure **S1**. Exchange and labeling efficiency of M2β-S1 A488RLC.

Representative SDS-PAGE and fluorescence imaging gel of M2β-S1 A488RLC used to examine exchange and labeling efficiency. M2β-S1 A488RLC was complexed with actin and spun down into a pellet, then released with excess ATP. The amount of A488RLC in the supernatant was quantified using a standard curve with a known concentration of A488RLC loaded on the gel. The total concentration of M2β-S1 A488RLC in the supernatant was determined by Bradford. Order of samples on the gel: 1, molecular weight marker; 2, M2β-S1 A488RLC protein before spin-down; 3, supernatant containing unbound A488RLC; 4, the supernatant of M2β-S1 A488RLC released from actin in the presence of ATP; 5, the final pellet of M2β-S1 A488RLC:actin complex; 6-9, standard loading curve of A488RLC; 10, M2β-S1 A488RLC protein; 11, M2β-S1 Non-Ex protein. The stoichiometry of A488 RLC to M2β-S1 was close to 1:1 (1:1 ± 0.2, for WT M2β-S1 A488RLC, $N = 3$).

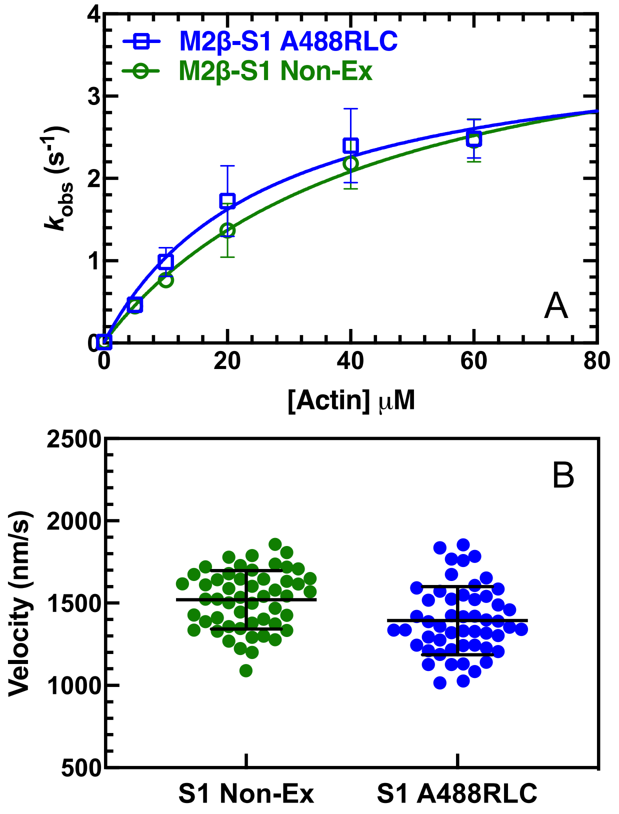


Figure **S2**. Steady-state ATPase and *in vitro* motility of non-exchanged and exchanged M2β-S1.
**(A)** The actin-activated ATPase activity of purified M2β-S1 A488RLC, and a non-exchanged control (M2β-S1 Non-ex) examined in parallel, was determined as a function of actin in MOPS 20 buffer at 25 ℃. The data points represent the average ± SD from 2 protein preparations. **(B)** The sliding velocity of M2β-S1 A488RLC and Non-Ex was measured in MOPS 20 buffer. 50 filaments in total at 0.4 μM loading concentration were analyzed manually, and velocities were pooled together to determine the average velocity. The black bar represents the mean ± SD. Data are summarized in Table 2.

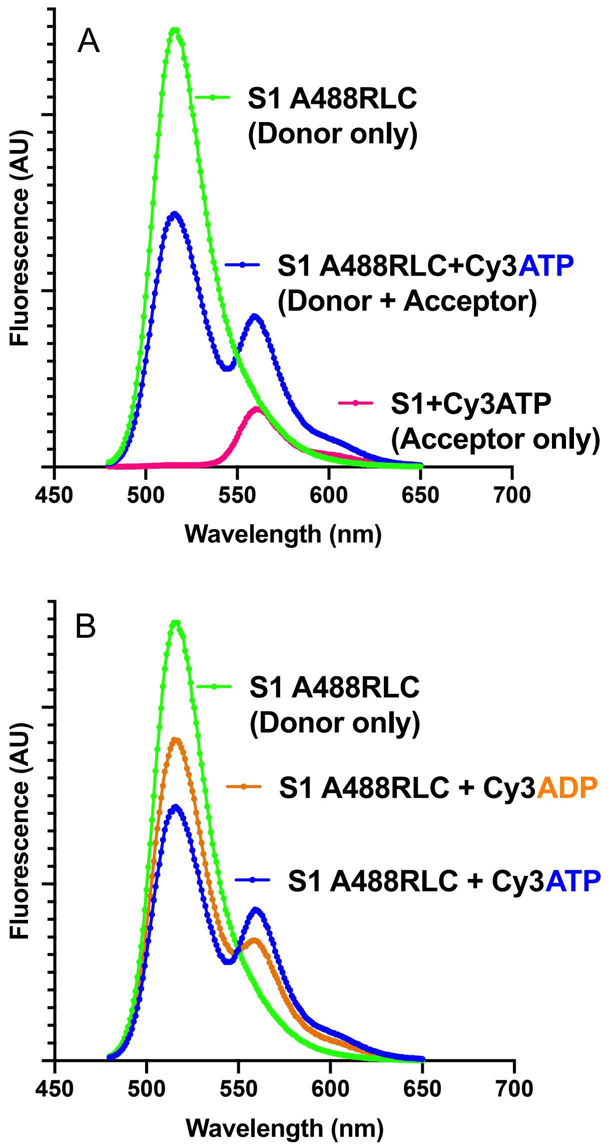


Figure S3. Examination of steady-state FRET as a function of nucleotide-state.
Representative wavelength scans used to perform FRET efficiency measurements of M2β-S1 A488 RLC with Cy3ATP and Cy3ADP. (A) The fluorescence spectrum of donor-acceptor, donor only, and acceptor only is shown. (B) The fluorescence spectrum of donor only compared with donor-acceptor and comparing Cy3ATP and Cy3ADP. The FRET efficiency was calculated with fraction of Cy3 nucleotide bound corrected, using the $K_D$ estimated from stopped flow measurements (Figure 1C). A summary of the FRET efficiency measurements and associated distances is shown in Table 1.

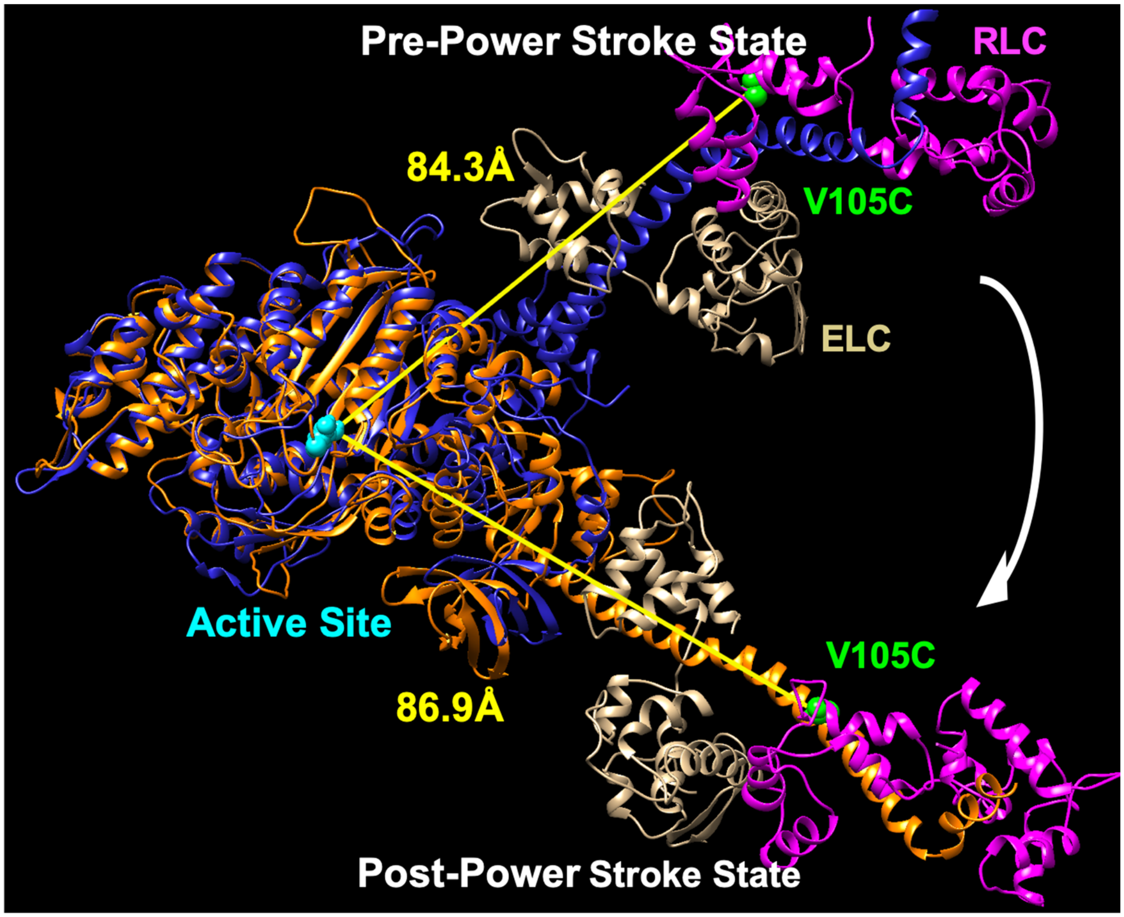


Figure **S4**. Structural alignment of M2β-S1 in pre- and post-power stroke conformations.

The pre- (blue) and post- power stroke (orange) homology structures of M2β-S1 were aligned using the structural alignment feature in the program Chimera. Essential light chain (ELC) is shown in tan, regulatory light chain (RLC) is shown in magenta, cysteine mutation introduced at residue 105 (V105C) in the RLC is shown in green, and the residue Asn-238 (N238) in the active site, which is proposed to be close to the Cy3 fluorophore, is shown in cyan. The distances between N238 and V105C were determined using the program Chimera. The M2β-S1 pre- and post-power stroke homology structures are acquired from the Spudich Lab website. (http://spudlab.stanford.edu/homology-models)

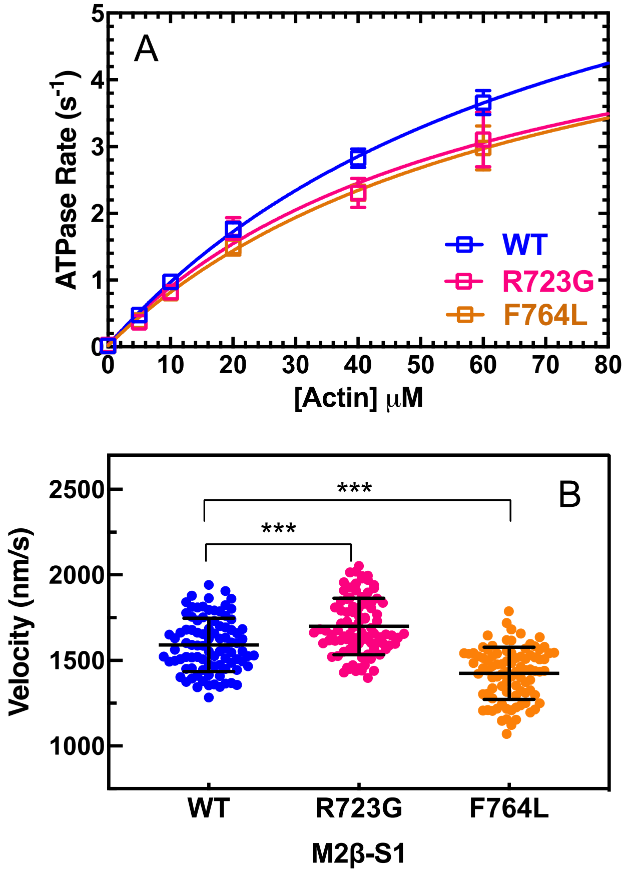


Figure **S5.** Steady-state ATPase and *in vitro* motility of M2β-S1 WT, R723G, and F764L.

**(A)** The actin-activated ATPase activity of purified M2β-S1 WT, R723G, and F764L was determined as a function of actin in MOPS 20 buffer at 25 ℃. The data points represent the average ± SEM from 3-6 protein preparations. **(B)** The sliding velocity of M2β-S1 WT, R723G, and F764L was measured in MOPS 20 buffer. 90 filaments in total (30 filaments from each protein preparation) at 0.24 μM loading concentration were analyzed manually, and velocities were pooled together to determine the average velocity. The black bar represents the mean ± SD. Unpaired student's *t*-tests were done to compare each mutant with WT (***, $p < 0.0001$).

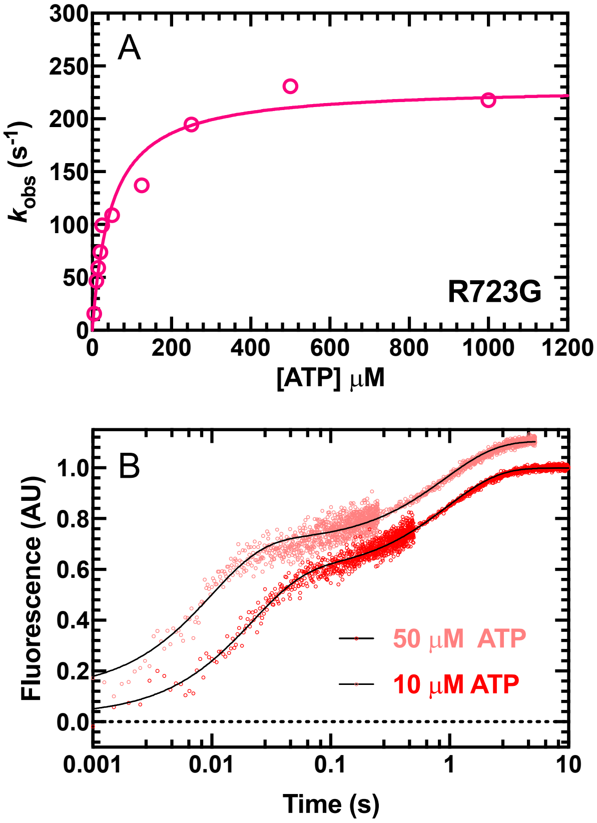


Figure **S6.** ATP binding and hydrolysis in M2β-S1 R723G.

Tryptophan fluorescence enhancement was used to monitor ATP binding and hydrolysis by mixing 1μM purified M2β-S1 R723G with varying concentrations of ATP (2.5-1000 μM). The fluorescence transients were best fit to a double exponential function. **(A)** The fast phase of the transients was plotted as a function of ATP concentration and fit to a hyperbolic function to determine the maximum rate of ATP hydrolysis and dependence on ATP concentration. Rate constants at low ATP concentrations (2.5-12.5 μM) were fit to a linear function to determine the second-order binding constant for ATP (see Table 2). **(B)** Representative fluorescence transients at 10 and 50 μM ATP fit to a double exponential function (10 μM ATP, $k_{Fast} = 47.0 \pm 0.9$ $s^{-1}$, $A_{Fast} = 0.58$; 50 μM ATP, $k_{Fast} = 102.2 \pm 2.4$ $s^{-1}$, $A_{Fast} = 0.60$).

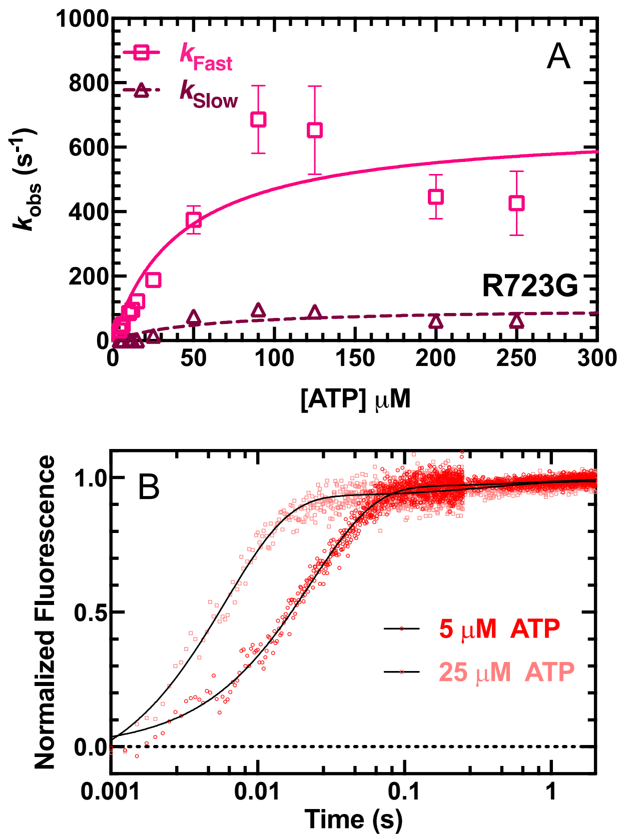


Figure **S7.** ATP binding to acto-M2β-S1 R723G.

ATP-induced dissociation from pyrene actin was performed by mixing a complex of M2β-S1 R723G:pyrene actin (0.375 μM M2β-S1 and pyrene actin) with varying concentrations of ATP (2 to 250 μM). The fluorescence transients were fit to a double exponential function. **(A)** Both fast and slow phase rate constants were plotted as a function of ATP concentration and fit to hyperbolic function to determine the maximum rate constant of the fast (668 ± 121 $s^{-1}$) and slow phase (102 ± 28 $s^{-1}$) and their dependence on ATP concentration. **(B)** Representative fluorescence transients at 10 and 25 μM ATP fit to a double exponential function (10 μM ATP, $k_{Fast}$ = 43.3 ± 0.4 $s^{-1}$, $A_{Fast}$ = 0.97; 25 μM ATP, $k_{Fast}$ = 166.7 ± 3.0 $s^{-1}$, $A_{Fast}$ = 0.95).

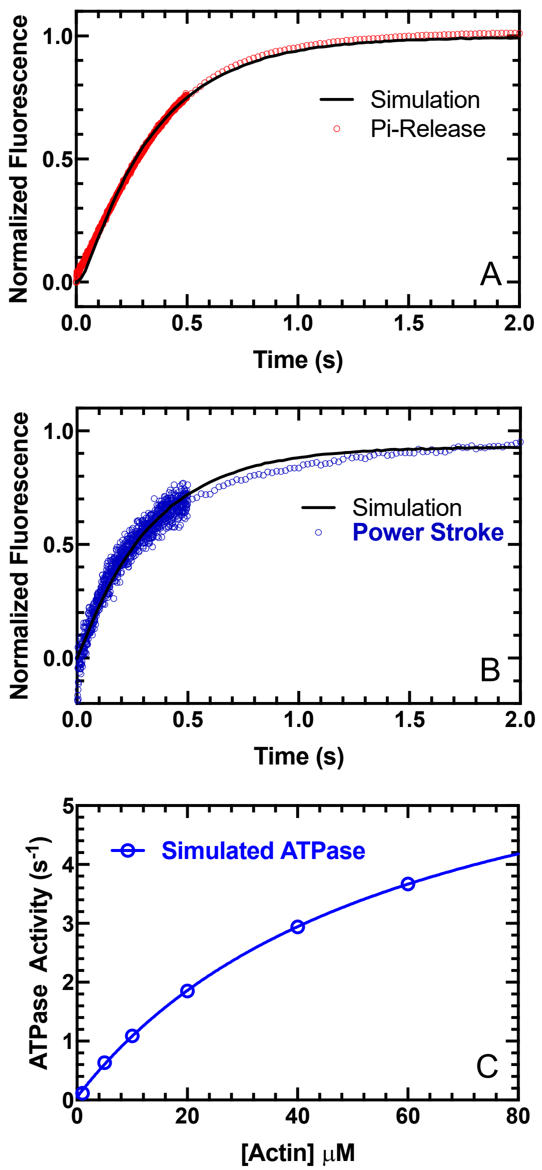


Figure **S8**. Experimental and simulated data.

**(A)** Representative experimental trace (red) of Pi release at 10 μM actin, fit to a double exponential function, is compared with the simulated trace of Pi release (black). **(B)** Representative experimental trace (blue) of the power stroke at 30 μM actin, fit to a double exponential function, is compared with the simulated trace of the power stroke using Kintek Explorer (black). **(C)** Simulated ATPase activity using the rate constants listed in Table S3. Simulations were performed in Kintek Explorer.

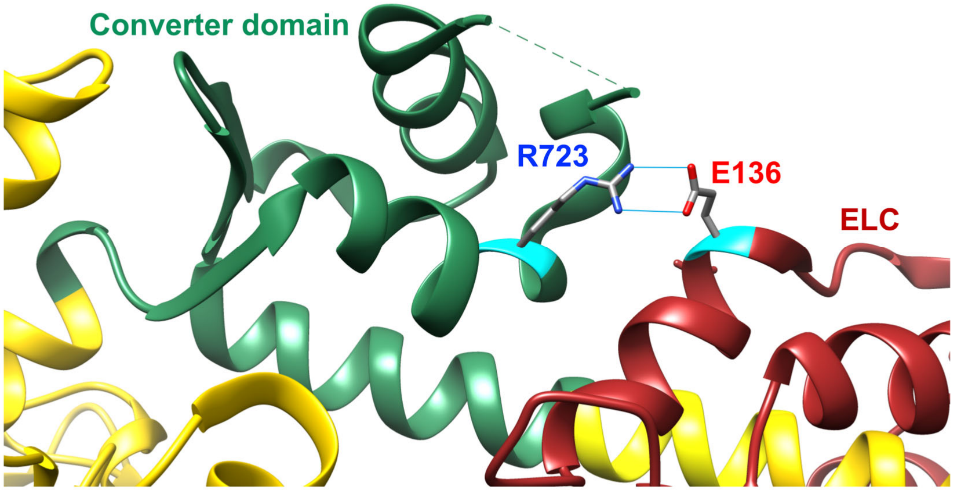


Figure **S9**. Structure of M2β-sS1 highlighting the Arg-723 residue.

Arg-723 (R723, blue) is located in the converter domain (green), close to the ELC (oxblood) associated with the lever arm. Arg-723 forms a salt-bridge with Glu-136 (E136, red) in the ELC. Structural modeling was done using the program Chimera. (PDB ID: 5N69)
